# The inactive X chromosome drives sex differences in microglial inflammatory activity in human glioblastoma

**DOI:** 10.1101/2024.06.06.597433

**Authors:** Marla E. Tharp, Claudia Z. Han, Maya Talukdar, Christopher D. Balak, Conor Fitzpatrick, Carolyn O’Connor, Sebastian Preissl, Justin Buchanan, Alexi Nott, Laure Escoubet, Konstantinos Mavrommatis, Mihir Gupta, Marc S. Schwartz, U Hoi Sang, Pamela S. Jones, Michael L. Levy, David D. Gonda, Sharona Ben-Haim, Joseph Ciacci, David Barba, Alexander Khalessi, Nicole G. Coufal, Clark C. Chen, Christopher K. Glass, David C. Page

## Abstract

Whether an individual is a biological female or male affects cancer risk, but the responsible mechanisms and cell types remain obscure. Glioblastoma multiforme (GBM) is a male-biased cancer that is highly aggressive, and resistant to treatment, with poor patient survival. Dismal prognoses in GBM are due in part to the specialized immune system of the brain, consisting largely of microglia, which regulate GBM development and progression. We hypothesized that microglia function differently in females and males and thereby contribute to the observed male bias in GBM. We sorted TAM-MGs (tumor-associated macrophages - microglia) from human GBMs and low-grade gliomas and performed bulk transcriptomic and epigenomic assays to identify sex-biased gene expression. We used published single-cell transcriptomic data from human GBMs to predict sex-biased TAM-MG interactions with other cell types. We found that female and male TAM-MGs mount different inflammatory responses, with female TAM-MGs displaying stronger interferon signaling and cytotoxic T-cell interactions that should enhance anti-tumor immunity in GBM. We validated these sex-differential inflammatory responses experimentally, and determined that genes on the sex chromosomes, specifically those expressed by Xi (the “inactive” X chromosome), drive these differences. Together, our results suggest that sex-differential TAM-MG inflammatory responses contribute to the higher incidence and mortality of GBM in males.

## Introduction

Glioblastoma multiforme (GBM) is the most common, most lethal primary brain tumor in adults^1^. GBM is highly resistant to treatment, with both 5-year survival and the standard of care unchanged since 2005^2,3^. Modern immunotherapies have not impacted GBM survival^4,5^. One reason GBM is so difficult to treat is the brain’s privileged immune environment, which is thought to be essential for neurological functions, but also supports tumor development and progression^6^. The blood-brain barrier (BBB) protects the brain from harmful substances and excessive inflammation, but also facilitates GBM development and resistance to treatment. Specifically, the BBB restricts the entry of circulating immune cells, including cytotoxic T-cells, which surveil and eliminate cancer cells in other tissues^7^. As a result, the immune landscape of brain tumors is dominated by microglia. Derived from the embryonic yolk sac, microglia are brain-resident macrophages that perform neurodevelopmental roles and mount immunological responses milder than those of blood-borne macrophages^8^. Second, BBB integrity is often disrupted during GBM development, permitting heterogeneous infiltration of circulating immune cells with diverse compositions, interactions, and phenotypes that make tumors difficult to treat with a single drug regimen^7,9,10^. Understanding how this specialized and dynamic brain immune environment impacts GBM tumor progression could lay the foundation for more effective treatments.

Biological sex influences GBM, with males showing increased incidence (male:female ratio 1.6:1) and higher mortality^1,11^. Since the immune system plays a vital role in controlling tumor development and progression^7^, and females typically display stronger immune responses^12^, we asked whether sex differences in tumor-immune interactions drive sex differences in GBM outcomes. Furthermore, increased immune cell infiltration is associated with the most male-biased GBM subtypes^13^, and may be involved in establishing sex differences. Given that microglia are the most abundant immune cells in the brain and heavily infiltrate GBM tumors, we hypothesized that sex differences in GBM stem from genetic and molecular mechanisms in TAM-MGs (tumor-associated macrophages - microglia)^14^.

Microglia display remarkable plasticity, constantly surveying and responding to alterations in their local environment, including tumors, by engaging sets of transcription factors that activate gene expression programs that in turn yield distinct phenotypic states^15,16^. The TAM-MG state is one of the more complex due to the evolving and heterogeneous tumor microenvironment^17^; the TAM-MG state can encompass both tumor-supportive and tumor-killing phenotypes^18–20^. Little is known about genetic and molecular mechanisms regulating TAM-MG phenotypes in human GBM, or how they might differ between males and females. As in humans, mouse glioma models display male-biased tumor growth and mortality, and recent studies suggest that these sex differences may be mediated by microglia. In one study, microglia-enriched expression of Junction Adhesion Molecule A (JAM-A) was found to regulate pathogenic immune activation exclusively in female tumors, leading to better survival outcomes in female mice^21^. In another mouse study, single-cell RNA sequencing of gliomas distinguishing subpopulations of tumor-associated myeloid cells revealed an increased interferon (IFN) response signature in female microglia and macrophages, while males showed an increased tumor supportive signature in macrophages only^8^. Since IFN signaling typically promotes anti-tumor effects, these results support the heightened ability of female (as compared with male) microglia to combat tumor cells.

To investigate sex differences in human TAM-MGs and their role in the male-biased incidence and mortality in GBM, we generated and analyzed transcriptomic and epigenomic data from FACS-isolated adult human TAM-MGs and control microglia. We utilized published single-cell RNA-seq (scRNA-seq) data of adult GBM tumors from newly diagnosed patients^22^ to validate sex differences observed in TAM-MGs and investigate their effects on other cell types in the GBM microenvironment. We found that, compared to males, female TAM-MGs exhibited stronger expression of anti-tumorigenic immune genes, especially those contributing to the type I IFN response, in low-grade gliomas and high-grade GBM. This sex difference may drive more efficient tumor cell killing in females through enhanced cytotoxic T-cell interactions. In contrast, male TAM-MGs showed stronger expression of pro-tumorigenic immune genes involved in NF-kB signaling that can enhance proliferation, angiogenesis, and immunosuppression, leading to worse GBM outcomes in males. Moreover, we demonstrated that sex differences in TAM-MG inflammatory responses are facilitated by genes expressed or modulated by the female-specific, so-called “inactive” X chromosome (Xi). Our studies support a pivotal role for TAM-MGs in establishing male-biased GBM incidence and mortality, and we directly link this male bias to the sex chromosomes.

## Results

### TAM-MGs exhibit reduced expression of genes involved in microglia maturation and anti-tumor immunity with increasing tumor grade

Before assessing sex differences in human TAM-MGs, we characterized the TAM-MG state through whole-genome analyses, both transcriptomic and epigenomic. To this end, we isolated TAM-MGs from adult brain tumor resections, including grade II and III gliomas and GBM (also known as grade IV) classified using genetic and morphological criteria established by the World Health Organization (Fig. 1A, Table S1)^23^. GBM is the highest-grade tumor and is distinguished from other gliomas by 1) wildtype isocitrate dehydrogenase *IDH* gene, 2) regions of necrosis, 3) excessive and aberrant neovascularization, 4) enhanced proliferation and spreading, and 5) increased macrophage infiltration^23,24^. As controls, we studied microglia isolated from non-epileptic portions of brain biopsies of individuals undergoing surgery for epilepsy^25,26^. TAM-MG and control microglia populations were FACS sorted using expression of CD11b^+^, CD45^mid^, CX3CR1^mid^, CD64^+^, and CCR2^lo^ to exclude inflammatory macrophages and recently migrated monocytes (Fig. 1B, Fig. S1A-D). We performed bulk RNA-seq, ATAC-seq, and H3K27ac ChIP-seq on sorted TAM-MGs and control microglia (Fig. 1B, Table S2).

**Figure 1.**
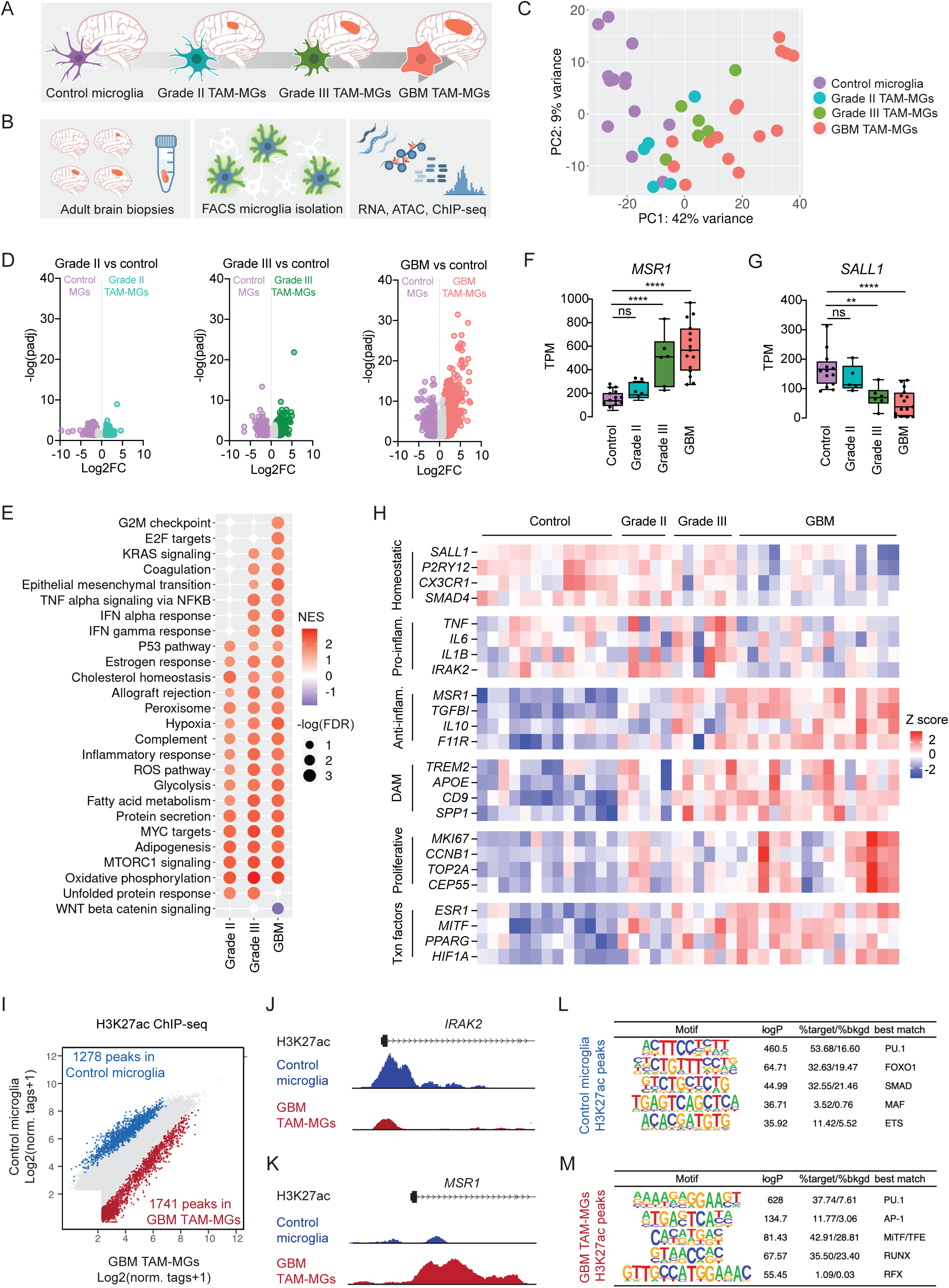
TAM-MGs exhibit reduced expression of genes involved in microglia maturation and anti-tumor immunity with increasing tumor grade. A. Overview of project design: investigating genetic and epigenetic regulators of human TAM-MG state using control microglia and TAM-MGs of grades II, III, and IV (GBM). B. Experimental design: isolation of microglia from human brain tumor resections and control brain tissue by FACS, followed by transcriptomic and epigenomic assays. C. Principal component analysis (PCA) of bulk RNA-seq libraries from human TAM-MGs and control microglia. D. Volcano plots of differentially expressed genes (DEGs) in TAM-MGs vs control microglia across tumor grades. Control microglia, n = 13; grade II TAM-MGs, n = 5; grade III TAM-MGs, n = 6; GBM TAM-MGs, n = 15. E. Enrichment of Hallmark pathway gene sets among grade II, III, and GBM TAM-MGs vs control microglia. NES = normalized enrichment score. FDR = false discovery rate. Only gene sets with - log (FDR) > 1.3 in one or more pairwise comparisons are listed. Gene sets with -log (FDR) > 1.3 are shown in color. F. Expression of *MSR1* correlates positively with tumor grade. G. Expression of *SALL1* correlates negatively with tumor grade. H. Genes representative of characterized microglia states that are differentially expressed in TAM-MGs compared to control microglia. Z score expression was calculated from log10 (TPM + 1) values for each gene. I. Differential H3K27ac ChIP-seq peaks (log2 (normalized tags +1)) in control microglia (blue) and GBM TAM-MGs (red). J. H3K27ac ChIP-seq data sets in the vicinity of the *IRAK2* gene locus from control microglia (blue) and GBM TAM-MGs (red). K. H3K27ac ChIP-seq data sets in the vicinity of the *MSR1* gene locus from control microglia (blue) and GBM TAM-MGs (red). L. Transcription factor binding motifs enriched in differential H3K27ac ChIP-seq peaks in control microglia. M. Transcription factor binding motifs enriched in differential H3K27ac ChIP-seq peaks in GBM TAM-MGs.

We performed principal component analysis (PCA) on bulk RNA-seq transcriptomes and observed that TAM-MG samples clustered primarily by tumor grade, and secondarily by other variables such as age, primary vs. recurrent tumor status, and *IDH* mutation status (Fig. 1C, Table S1). To describe the TAM-MG state in our samples, we identified differentially expressed genes (DEGs) between control microglia and TAM-MGs of each grade individually (Fig. 1D, Table S3). We found that TAM-MGs in GBM had greater numbers of DEGs than those in lower-grade gliomas, and that many of those DEGs were unique to GBM TAM-MGs, underscoring substantial differences between grades in the tumor microenvironment. We performed gene set enrichment analysis (GSEA) on the control vs TAM-MG comparisons for each tumor grade to investigate how TAM-MG pathways were affected by these different tumor environments (Fig. 1E). Querying the fifty “Hallmark” gene sets^27^, we observed that some sets were enriched in TAM-MGs from all tumor grades compared to control microglia, with other gene sets enriched only in TAM-MGs from high grades (Fig. 1E). Gene sets enriched in TAM-MGs from all grades included metabolic processes, upregulation of which may be required for TAM-MG survival in the tumor microenvironment (Fig. 1E). In TAM-MGs from grade III and GBM tumors, we observed enrichment of gene sets involved in inflammation, including IFN responses and TNF alpha signaling via NFKB (Fig. 1E). Such inflammatory programs can drive both acute anti-tumor responses, or if persisting, chronic pro-tumor responses^28^. Last, in TAM-MGs from GBM specifically, we observed enrichment of proliferative pathways (Fig. 1E). GBMs exhibit a substantially increased proliferation rate compared to gliomas, indicating an environment abundant in mitogens that may influence TAM-MGs^20^.

We next explored genes associated with known microglial phenotypic states, and how their expression changed with advancing tumor grade. Among the genes whose expression increased with tumor grade was the known TAM-MG biomarker *MSR1/CD204*, associated with immunosuppression and decreased survival in GBM (Fig. 1F)^29^. By contrast, *SALL1*, a microglial lineage-determining and homeostatic gene, decreased with tumor grade (Fig. 1G)^30^. Similarly, expression of the microglia homeostatic genes *P2RY12* and *CX3CR1* declined with tumor grade, while expression of the immunosuppressive genes *IL10* and *F11R (JAM-A)* increased with tumor grade (Fig. 1H). Interestingly, expression of the pro-inflammatory genes *IL6* and *TNF* increased in grade II-III gliomas compared to control microglia, but decreased in GBM, highlighting the dynamic nature of TAM-MG gene expression during tumor progression (Fig. 1H). Expression of genes previously identified as determinants of the disease-associated microglia (DAM) state in neurodegenerative disease^31^, and proliferating microglia, increased in expression with tumor grade in TAM-MGs (Fig. 1H).

The transcription factors *MITF*, *PPARG*, *ESR1*, and *HIF1A* were expressed more abundantly in higher-grade tumors (Fig. 1H), potentially regulating the TAM-MG state. To test how the TAM-MG state is regulated, we assayed changes in microglia enhancer activation in GBM tumors. Previous studies have demonstrated that microglial phenotypes in disease are governed by transcription factor binding and enhancer activation in response to local environmental cues^16^. We performed H3K27ac ChIP-seq on the TAM-MGs to identify active promoters and enhancers, and we specifically identified enhancers whose activity differs between human GBM TAM-MGs and control microglia. Comparison of the active enhancer landscape in GBM TAM-MGs and control microglia revealed 1741 regions more active in GBM TAM-MGs and 1278 regions more active in control microglia (Fig. 1I). For example, H3K27ac signal in the promoter of *MSR1* was greater in GBM TAM-MGs than in control microglia, while the reverse was true for *IRAK2*, supporting the differences in transcription that we observed for these genes (Fig. 1J-K). Application of *de novo* motif analysis showed enrichment for motifs for transcription factors including the SMAD family members in differential H3K27ac peaks in control microglia and the MiTF-TFE family members in differential H3K27ac peaks in GBM TAM-MGs (Fig. 1L-M). SMAD4 interacts with SALL1 to promote microglia maturation during fetal brain development^30^, while MiTF-TFE factors are master regulators of lysosomal function, autophagy, and phagocytosis^32^. Since *SMAD4* and other microglia maturation genes decrease in expression in TAM-MGs with increasing tumor grade, while *MITF* along with other phagocytic genes in the DAM state increase in expression in TAM-MGs with increasing tumor grade, these families of transcription factors may regulate the TAM-MG state.

Cancer cells sometimes assume the properties of their progenitors, leading to increased invasiveness, immune evasion, and drug resistance^33^. Since we observed that TAM-MGs exhibit reduced expression and accessibility of genes involved in microglia maturation, we asked whether tumor association drives microglia toward a progenitor, fetal-like state. We compared GBM TAM-MG transcription factor families to those recently identified in human microglia along a developmental context, specifically, postnatal compared to fetal microglia^25^. SMAD motifs, which were enriched in control microglia compared to GBM TAM-MGs, were also enriched in postnatal microglia compared to fetal microglia. MiTF-TFE motifs, which were enriched in GBM TAM-MGs compared to control microglia, were also enriched in fetal microglia compared to postnatal microglia^26^. *MITF* expression that increases in TAM-MGs with tumor grade, also is increased in fetal compared to postanal microglia (Fig. S2A). In contrast, *SMAD4* expression that decreases in TAM-MGs with tumor grade is also decreased in fetal compared to postnatal microglia (Fig. S2B)^26^. Further, we compared all DEGs between TAM-MGs vs. control microglia and fetal vs. postnatal microglia, and found a significant overlap in the identity and directionality of DEGs between the two comparisons (Fig. S2C). Collectively, these results suggest that the GBM tumor microenvironment influences human microglia to assume a more fetal state that potentiates their development as opposed to immune regulation, supporting tumor growth.

### Female and male TAM-MGs mount different responses in low-grade gliomas and GBM

Given the male-biased incidence and mortality rate in GBM, and precedence for sex differences in immune regulation, we asked if TAM-MGs display sex-biased gene expression that may explain these differences. We analyzed sex-biased genes in TAM-MGs from pooled grade II and grade III gliomas, and from GBM, as well as from control microglia to determine whether sex differences observed in TAM-MGs are established in the homeostatic state (Fig. 2A-C, Table S4A-B).

**Figure 2.**
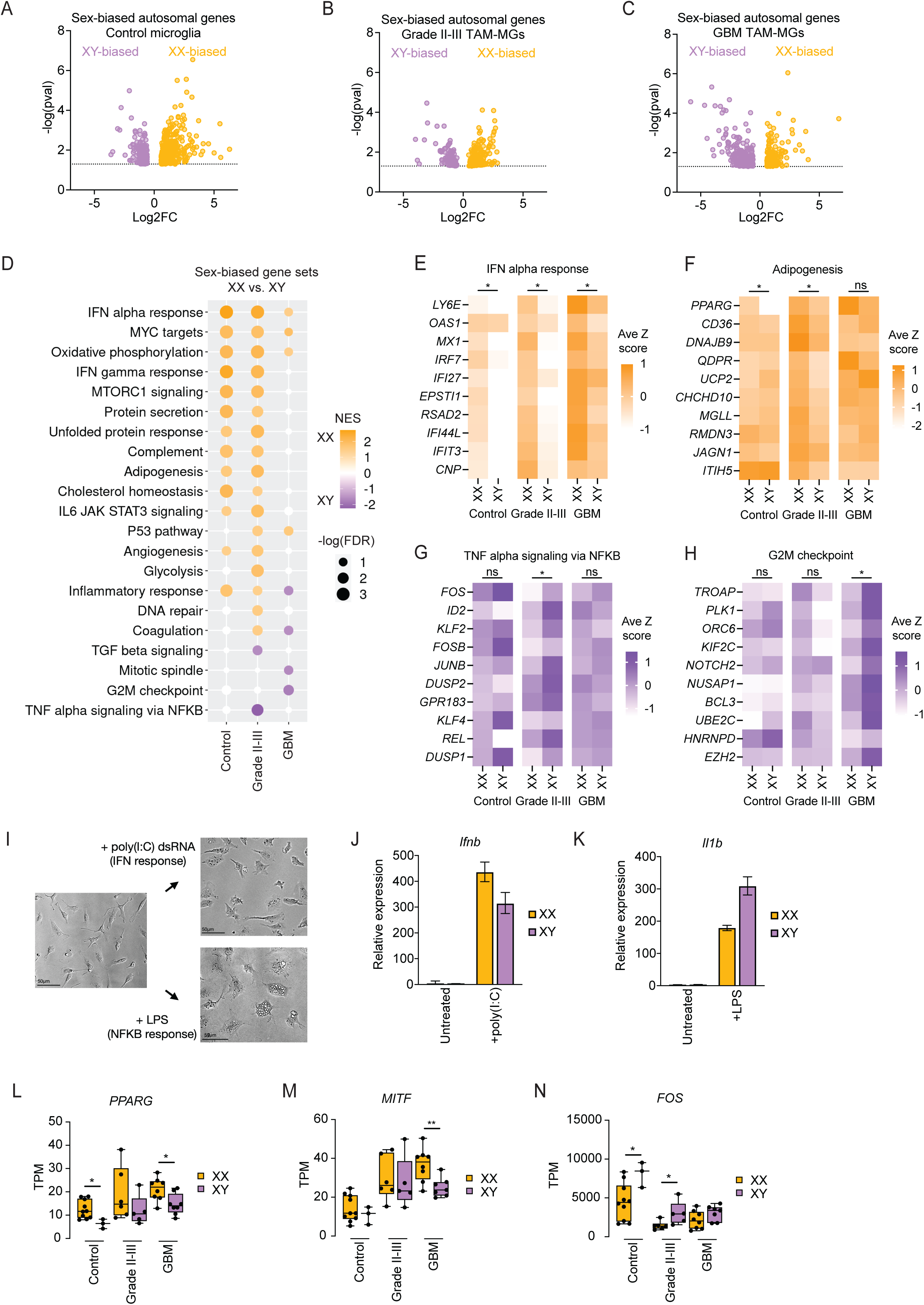
TAM-MGs mount sex-biased responses in low-grade gliomas and GBM. A. Sex-biased autosomal genes in control microglia. XX, n = 10; XY, n = 3. B. Sex-biased autosomal genes in grade II-III TAM-MGs. XX, n = 6; XY, n = 5. C. Sex-biased autosomal genes in GBM TAM-MGs. XX, n = 8; XY, n = 7. D. Enrichment of Hallmark pathway gene sets among sex-biased genes in control microglia, grade II-III TAM-MGs, and GBM TAM-MGs. NES = normalized enrichment score. FDR = false discovery rate. Only gene sets with -log (FDR) > 1.3 in one or more pairwise comparisons are listed. Gene sets with -log (FDR) > 1.3 are shown in color. E. Top 10 leading-edge genes of XX-biased Interferon alpha response gene set in grade II-III TAM-MGs. F. Top 10 leading-edge genes of XX-biased Adipogenesis gene set in grade II-III TAM-MGs. G. Top 10 leading-edge genes of XY-biased TNF-alpha-signaling-via-NFKB gene set in grade II-III TAM-MGs. H. Top 10 leading-edge genes of XY-biased G2M checkpoint gene set in GBM TAM-MGs. I. Brightfield images of FACS-isolated mouse microglia cultured in vitro for three days, treated with poly(I:C) for 24 h, and treated with LPS for 24 h. Scalebar = 50 µm. J. Expression of *Ifnb* in XX and XY microglia after poly(I:C) treatment for 4 h. K. Expression of *Il1b* in XX and XY microglia after LPS treatment for 4 h. L. Expression of *PPARG* in XX and XY control microglia and TAM-MGs. Significantly XX-biased in control microglia and GBM TAM-MGs. M. Expression of *MITF* in XX and XY control microglia and TAM-MGs. Significantly XX-biased in GBM TAM-MGs. N. Expression of *FOS* in XX and XY control microglia and TAM-MGs. Significantly XY-biased in control microglia and grade II-III TAM-MGs.

We performed GSEA on sex-biased genes from control microglia, grade II-III TAM-MGs, and GBM TAM-MGs, and found that female and male TAM-MGs mount different immune responses. First, grade II-III TAM-MGs showed female enrichment of pathways related to anti-tumor inflammatory activity (IFN alpha response, IFN gamma response, and IL6 JAK STAT3 signaling), as well as pathways involved in lipid metabolism (adipogenesis and cholesterol homeostasis) (Fig. 2D-F). In contrast, male enrichment of pathways involved in tumor-supportive inflammatory activity (TGF beta signaling and TNF alpha signaling via NFKB) were observed in grade II-III TAM-MGs (Fig. 2D, G). In GBM TAM-MGs, the IFN alpha response was again enriched in female samples, this time along with the p53 pathway (Fig. 2D, E). Male GBM TAM-MGs were enriched in proliferative pathways (G2M checkpoint and mitotic spindle) (Fig. 2D, H). We performed GSEA querying gene sets induced by interferon gamma treatment in cultured human fetal microglia^34^ to validate female-biased IFN signaling in TAM-MGs (Fig. S3A-B). Control microglia showed female enrichment of IFN responses and lipid metabolism gene sets, similar to grade II-III TAM-MGs, suggesting that these sex differences may be established in the homeostatic state and enhanced with tumor association (Fig. 2D-F).

We experimentally validated sex-biased immune responses using male and female mouse microglia isolated by FACS and cultured in vitro. We stimulated microglia with poly(I:C) to activate IFN signaling and lipopolysaccharide (LPS) to activate NF-kB signaling (Fig. 2I). We found that female microglia mounted a stronger response to poly(I:C) based on induction of IFN-stimulated gene *Ifnb* (Fig. 2J). In contrast, male microglia mounted a stronger response to LPS based on induction of the pro-inflammatory gene *Il1b* (Fig. 2K). These results validate the sex differences in immune responses observed in human TAM-MGs, where female TAM-MGs are enriched for IFN signaling genes and male TAM-MGs are enriched for TNF alpha signaling via NFKB genes.

We then asked how sex differences in TAM-MGs are regulated. We several transcription factors with sex-biased expression in TAM-MGs, including female-biased *PPARG* and *MITF* (Fig. 2L-M), and male-biased *FOS* expression (Fig. 2N). *PPARG* is a member of the female-biased adipogenesis gene set and a known anti-inflammatory and anti-tumor factor that may contribute to better GBM outcomes in females^35,36^. Conversely, FOS is a member of the male-biased TNF alpha signaling via NFKB gene set and linked to poor survival in malignant glioma and GBM patients that may contribute to worse GBM outcomes in males^37^.

Together, our analysis of autosomal sex-biased genes in TAM-MGs and control microglia suggest a greater ability of female TAM-MGs to mount an acute IFN response that suppresses tumor growth. Male TAM-MGs express more immunosuppressive and proliferative genes that support tumor growth and worsen GBM outcomes.

### Female TAM-MGs display enhanced interactions with cytotoxic T-cells

We next re-analyzed previously published scRNA-seq data from newly diagnosed GBM tumor resections to ask 1) whether the sex-biased inflammatory pathways observed in our human bulk sorted TAM-MGs could be recapitulated in an independent dataset, and 2) whether TAM-MG interactions with other immune and tumor cells in the GBM microenvironment are involved in these sex differences^22^. We analyzed 5 female and 6 male samples to find sex-biased gene expression in each major cell population in the GBM tumor microenvironment. Using enrichment of known marker genes, we subsetted four major cell types: TAM-MGs, tumor-associated bone marrow-derived macrophages (TAM-BMDMs), T-cells, and tumor cells (Fig. 3A, Fig. S4A-D). Next, we identified sex-biased genes for each of these four cell types, and determined which were unique versus shared between the cell types. T-cells and TAM-MGs had the most sex-biased genes, while tumor cells had the least, suggesting that immune cell populations establish sex differences (Fig. 3B). We went on to perform GSEA to determine sex-biased Hallmark gene sets for each of the cell types. We found that sex differences in the computationally subsetted GBM TAM-MGs were similar to those observed in our bulk-sorted TAM-MGs, with IFN alpha responses appearing the most XX-biased, suggesting female-biased anti-tumor immune effects through this pathway (Fig. 3C). Also consistent with bulk-sorted TAM-MGs from low-grade gliomas, TNF alpha signaling via NFKB was the most XY-biased gene set in TAM-MGs in the scRNA-seq (Fig. 3C).

**Figure 3.**
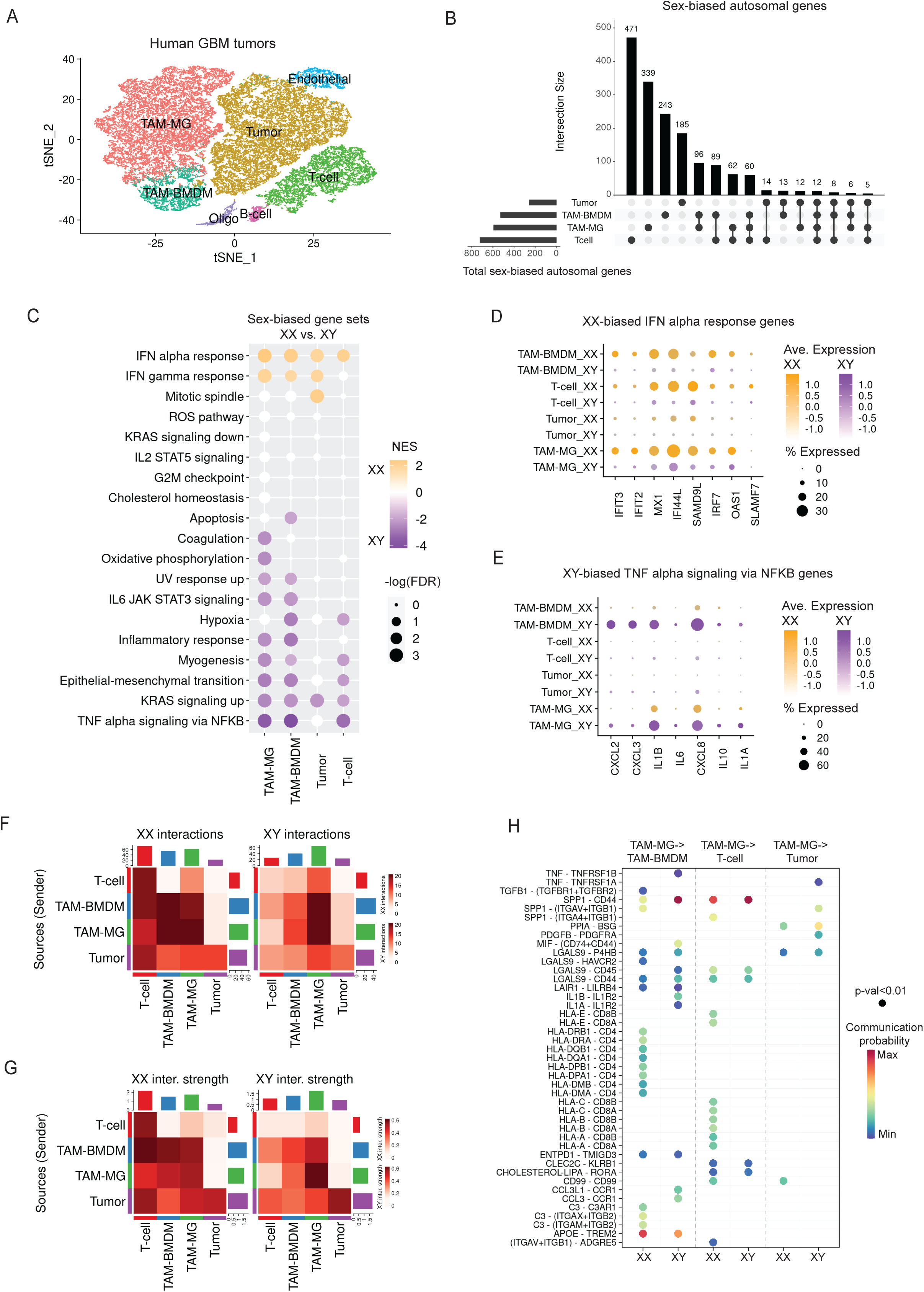
Female TAM-MGs display enhanced interactions with cytotoxic T-cells. A. Clustering of scRNA-seq data using Seurat from 11 adult human GBM samples, 5 XX and 6 XY, including TAM-MGs, TAM-BMDMs, T-cells, tumor cells, endothelial cells, B-cells, and oligodendrocytes. B. Number of sex-biased autosomal genes unique and shared between four major cell types: TAM-MGs, TAM-BMDMs, T-cells, and tumor cells. C. GSEA of sex-biased genes in four major cell types considering Hallmark categories. Significant XX-biased genes sets shown in orange and XY-biased in purple. D. Leading edge genes for the IFN alpha response in four major cell types. Plot shows percent of cells expressing respective gene. XX sample expression indicated in orange, XY in purple, with color intensity reflecting greater average expression. Genes plotted are also XX-biased in bulk sorted TAM-MGs. E. Leading edge genes for the TNF alpha signaling via NFKB response in four major cell types. Plot shows percent of cells expressing respective gene. XX sample expression indicated in orange, XY in purple, with color intensity reflecting greater average expression. Genes plotted are also XY-biased in bulk sorted TAM-MGs. F. Number of predicted receptor-ligand interactions between four cell types in XX and XY samples. G. Interaction strengths of predicted receptor-ligand interactions between four major cell types in XX and XY samples. H. Probability and significance of individual receptor-ligand interactions between TAM-MGs -> TAM-BMDMs, TAM-MGs -> T-cells, and TAM-MGs -> tumor cells in XX and XY samples.

Interestingly, non-TAM-MG cell types also showed significant XX-biased expression of IFN alpha response genes and XY-biased expression of TNF alpha signaling via NFKB genes (Fig. 3C). Because of this, we hypothesized that these sex-biased immune responses may involve interactions between TAM-MGs and other cell types. To further explore this cross-talk, we found expression of gene driving enrichment of the IFN alpha signaling and TNF alpha signaling via NFKB gene sets across cell types. The cell types with strongest XX-biased expression of IFN alpha signaling driver genes were in TAM and T-cell populations (Fig. 3D), while the strongest XY-biased expression of TNF alpha signaling via NFKB driver genes were in TAM populations only (Fig. 3E). Interestingly, tumor cells showed negligible expression of these same genes for either sex-biased response, again emphasizing the critical role of immune cells in sex differences in GBM (Fig. 3D-E). To test whether XX-biased IFN responses affect these anti-tumor cell-cell interactions, we utilized CellChat, which infers cell-cell communication networks from scRNA-seq data based on the expression of genes involved in known signaling pathways and receptor-ligand pairs^38^. We measured the number and strength of interactions between TAM-MGs and the other three cell types in XX and XY samples, and found that XX GBM tumors contained more and stronger interactions between TAM-MGs and T-cells, as well as TAM-MGs and TAM-BMDMs (Fig. 3F-G). In contrast, XY GBM tumors displayed more and stronger TAM-MG-to-TAM-MG interactions (Fig. 3F-G). Neither sex showed strong interactions between TAM-MGs and tumor cells (Fig. 3F-G). Teasing apart the individual receptor-ligand interactions driving these sex differences, we found that the XX-biased TAM-MG-to-T-cell interactions were primarily due to TAM-MG HLA to T-cell CD8 signaling that is cytotoxic and confers anti-tumor effects (Fig. 3H). Additionally, TAM-MG HLA to TAM-BMDM CD4 signaling was a prominent XX-biased interaction that supports greater activation of TAM-BMDMs in XX GBM tumors (Fig. 3H). These sex-biased interactions support the literature that IFN signaling in GBM involves crosstalk between TAM-MGs and T-cells, and this sensitizes tumor cells to CD8+ T-cell and TAM-MG phagocytosis-mediated killing^39^. Overall, our scRNA-seq analysis of XX and XY GBM tumors provides validation and a more comprehensive understanding of XX-biased IFN responses in TAM-MGs and their anti-tumor effects through T-cell and other GBM immune cell interactions.

### Sex-biased immune responses in TAM-MGs are regulated by the Xi

We proceeded to investigate the genetic and molecular basis of sex-biased immune responses in TAM-MGs, specifically, the roles of the sex chromosomes. Since the sex chromosomes are the genetic foundation of sex differences, we hypothesized that they encode drivers of sex-biased gene expression observed in TAM-MGs. Subsets of genes on the sex chromosomes are strong candidates based on their evolutionary histories and dosage sensitivities (Fig. 4A). Specifically, the X chromosome comes in two epigenetically distinct forms, the active (Xa) and inactive (Xi) forms, to account for dosage differences between XX and XY individuals. However, in humans, Xi maintains the expression of about one-third of its genes, although at attenuated levels, leading to their increased expression in females compared to males^40^. X chromosome genes that have retained homologs on the Y chromosome typically exhibit the highest and most cell-type-conserved expression from Xi, since Y homologs can compensate for some dosage differences^41^. However, X-Y pairs often diverge in sequence, expression, and function due to absence of genetic recombination between the X and Y chromosomes, which may facilitate sex differences^42^. Last, the Xi can modulate genes from Xa in trans, both positively and negatively, leading to sex-biased expression that is typically cell-type and context-specific^40^.

**Figure 4.**
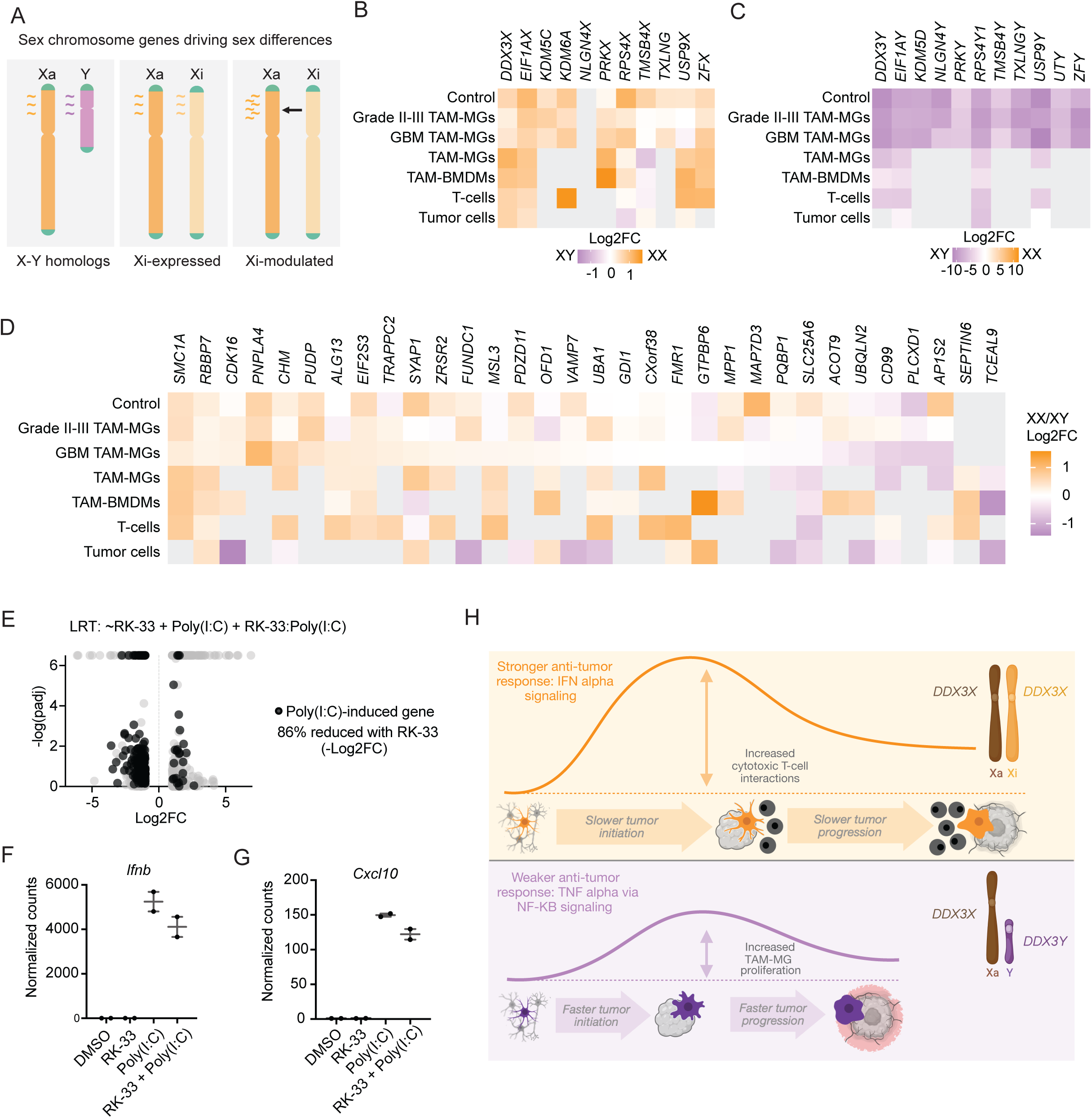
Sex-biased immune responses in TAM-MGs are regulated by the Xi. A. Diagram of sex chromosome encoded genes with potential to drive sex differences in TAM-MGs B. Sex-biased expression of Xi-expressed genes with Y homologs in control microglia, grade II-II TAM-MGs, GBM TAM-MGs, and subsetted GBM TAM-MGs, TAM-BMDMs, T-cells, and tumor cells. C. Sex-biased expression of Y-expressed genes in control microglia, grade II-II TAM-MGs, GBM TAM-MGs, and subsetted GBM TAM-MGs, TAM-BMDMs, T-cells, and tumor cells. D. Sex-biased expression of Xi-expressed and Xi-modulated genes without Y homologs in control microglia, grade II-II TAM-MGs, GBM TAM-MGs, and subsetted GBM TAM-MGs, TAM-BMDMs, T-cells, and tumor cells. E. Genes significantly affected by the interaction between 5uM RK-33 and 10uM poly(I:C) in HMC3 microglia. Log2FC > 1 or < -1. -log(padj) > 1.3. n = 2 for each condition. Genes significantly upregulated with poly(I:C) stimulation in control HMC3 microglia shown in pink. F. Expression of *Ifnb1* in mouse primary microglia treated with DMSO, 5uM RK-33, 10uM poly(I:C), or a combination. n = 2 for each condition. G. Expression of *Cxcl10* in mouse primary microglia treated with DMSO, 5uM RK-33, 10uM poly(I:C), or a combination. n = 2 for each condition. H. Model for sex differences in GBM driven by TAM-MGs, the sex chromosomes, and sex hormones.

To interrogate the role of the sex chromosomes in facilitating sex-biased immune responses in TAM-MGs, we first quantified sex-biased expression of Xi-expressed, Xi-modulated, and Y-expressed genes, as determined in human fibroblasts and lymphoblastoid cell lines with sex chromosome anueploidies^40^. In grade II-III and GBM TAM-MGs, control microglia, and the four subsetted GBM cell types, we observed Xi-expressed genes with Y homologs significantly female-biased, and Y-expressed genes significantly male-biased (Fig. 4B-C). We then queried sex-biased expression of Xi-expressed and Xi-modulated genes that do not possess Y homologs, and found that many also show female-biased expression, although more variable results across cell types than Xi-expressed genes with Y homologs (Fig. 4D).

Among the Xi-expressed genes with Y homologs showing female-biased expression across GBM cell types was the DEAD box helicase *DDX3X. DDX3X* has been implicated in IFN signaling^43,44^ and shows functional differences^45^ and compensatory relationships^46^ with its Y homolog, *DDX3Y* that make it a strong candidate driver of sex-biased inflammatory responses in TAM-MGs. Specifically, due to differences in the N-terminal domain between DDX3X and DDX3Y, DDX3Y is more readily sequestered into stress granules whereas DDX3X remains cytosolic and accessible to participate in innate immune responses, which we hypothesize contributes to stronger anti-tumor immune responses in female TAM-MGs^45^.

To examine the role of *DDX3X* in the female-biased IFN response in microglia, we treated the human microglia cell line with the small molecule inhibitor RK-33, prior to stimulation with poly(I:C)^47^. RK-33 inhibits ATPase activity of DDX3, and inhibits cell cycle progression in cancer cells^47^. We observed that RK-33 also inhibited cell cycle activity in HMC3 microglia (Fig. S5A-B). When stimulated with poly(I:C), we observed that RK-33 treatment resulted in the dampening of the poly(I:C) induced genes in HMC3 microglia, which share a large overlap with XX-biased IFN response genes in TAM-MGs (Fig. S5C-D, Fig. 4E). We validated this result using non-proliferative primary mouse microglia, and indeed, inhibition of DDX3X with RK-33 resulted in reduced expression of poly(I:C)-induced genes, including *IFNB1* and *CXCL10* by poly(I:C) (Fig. 4F-G).

Collectively, our results support that sex-biased immune responses in human TAM-MGs involve contributions from the sex chromosomes, particularly Xi-expressed genes, which drive a stronger IFN response observed in female TAM-MGs. We demonstrated that *DDX3X*, a female-biased gene in TAM-MGs and Xi-expressed in a number of human tissues, is a key modulator of IFN-stimulated genes, thereby representing a candidate genetic driver of enhanced anti-tumor immunity and better GBM outcomes in females.

## Discussion

### A proposed model: sex-biased responses in TAM-MGs drive sex differences in GBM

We investigated male and female TAM-MGs from human brain malignancies to derive a model of the genetic molecular mechanisms of male-bias in GBM. By integrating gene expression data from sorted TAM-MG bulk RNA-seq and computationally subset TAM-MG single-cell RNA-seq, we make significant advances in asserting TAM-MGs as the cell type establishing male-bias in GBM through sex-biased anti-tumor immune responses. Using human and mouse microglia model systems, we demonstrate that the sex chromosomes, specifically, genes expressed by the inactive X chromosome, as the genetic basis of these sex-biased immune responses originating in TAM-MGs.

First, we analyzed sex differences in gene expression in sorted human TAM-MGs from both low-grade gliomas and GBM and found sex-biased immune responses that were consistent in tumors of both grades, as well as in an independent human TAM-MG population from GBM scRNA-seq data^22^. Specifically, female TAM-MGs showed enrichment of genes involved in type I IFN signaling, while male TAM-MGs were enriched for genes involved in NF-kB signaling (Fig. 4H). These immune responses can lead to different types of cellular interactions in the GBM tumor microenvironment, including enhanced interactions with cytotoxic T-cells through IFN signaling in females^39^, and enhanced immunosuppression through myeloid-derived suppressor cells through NF-kB signaling in males^48^. Further, transition to the more aggressive and mal-biased mesenchymal GBM subtype occurs in a NF-kB-dependent manner^49^. Pro-inflammatory and proliferative microglia have also been associated with high-grade GBM and may reflect chronic inflammation that promotes tumor growth^20^. We observed a male bias in these pro-inflammatory and proliferative pathways in GBM TAM-MGs that may contribute to worse GBM outcomes.

Finally, we show that Xi-expressed genes contribute to sex differences in TAM-MG immune responses. Xi-expressed and modulated genes have been quantified using aneuploidy cells in both in vitro-cultured fibroblasts and LCLs, as well as in vivo-isolated CD4+ T cells and monocytes^40,41^. We quantified the expression of these Xi-expressed and modulated genes in TAM-MGs to identify candidate genetic drivers of sex-biased immune responses. We tested the role of Xi-expressed, female-biased gene *DDX3X* in the microglia response to poly(I:C) as a proxy for the XX-biased IFN response in TAM-MGs using human and mouse microglia models. Our findings implicate *DDX3X* as an Xi-expressed driver of female-biased anti-tumorigenic IFN activity in TAM-MGs, and thus, better GBM outcomes in females compared to males (Fig. 4H). *DDX3X* has known roles in promoting the IFN response^43,44^ and a less functional Y homolog^45^. For example, *DDX3X* and *DDX3Y* diverge in the 5’UTR, which upon stress, results in the preferential sequestration of *DDX3Y* into stress granules rather than participating in inflammasome activity like *DDX3X*^45,50^. We speculate that the functionally divergent Y homologs and female-biased expression of X homologs of these X-Y pairs may be drivers of sex differences in TAM-MGs influencing inflammatory activity and GBM progression.

### Sex hormone interactions influence sex-biased TAM-MG immune responses

Although this study focused on the genetic drivers of sex differences from the sex chromosomes, an IFN – NF-kB axis has been described in individuals undergoing gender affirming hormone therapy, where testosterone treatment increases expression of NF-kB and decreases expression of IFN signaling genes in whole blood^51^. Given that IFN signaling genes are XX-biased and NF-kB signaling genes are XY-biased in our TAM-MG data, this same axis may be influenced by respective male and female sex hormone milieu (Fig. 4H).

### Loss of microglia maturation accompanies the TAM-MG state

We delineated the overall homeostatic microglia-to-TAM-MG transcriptomic landscape in both sexes and found that TAM-MGs in low-grade gliomas retain more features of mature microglia and anti-tumorigenic immune activity than do TAM-MGs in GBM. For example, we observed more pro-inflammatory gene expression in TAM-MGs from grade II-III gliomas compared to GBM TAM-MGs, which expressed higher levels of anti-inflammatory genes and additional pro-tumorigenic pathways such as those involved in cell proliferation. Using H3K27ac ChIP-seq in control microglia compared to GBM TAM-MGs, we showed that this reversion of microglia maturation in the TAM-MG state is controlled by epigenetic rewiring of active enhancer elements. We found that predicted transcription factor binding motifs enriched in GBM TAM-MGs were similar to those enriched in fetal microglia, supportive of microglia losing features of maturation in the GBM tumor microenvironment.

### Sex differences and advances in GBM immunotherapies

Clinical trials for immune checkpoint inhibitors in GBM have shown better results in males^52^. Our work supports that this sex difference may be due to males benefiting from therapeutically-enhanced cytotoxic T-cell interactions more than females, who normally have better T-cell effector function^53^. Combination treatment of immune checkpoint inhibitors with oncolytic viral immunotherapies are also under development^54^. Our work supports that sex-biased responses to this type of therapy is an important consideration, since females tend to mount stronger responses to viral infection^12^, and this may influence GBM progression and efficacy of the treatment differently than in males.

### Sex-biased microglia immune responses impact sex differences in neurological disorders

Neuroinflammation is a common process in the progression of neurological disorders, many of which also show sex differences. For example, autism shows a striking 4:1 male:female bias^55^ and Alzheimer’s disease a 1:2 male:female bias^56^. Our study of sex-biased immune responses in TAM-MGs establishing sex differences in GBM raises the question of whether microglia are the sex-biasing cell type in other neurological disorders that manifest differently in males and females. Evidence in support of this is that Alzheimer’s disease is exacerbated by neuroinflammation, and the predisposition of female TAM-MGs to mount a stronger IFN response may contribute to the female-biased incidence of this disease^57^.

### Key Resources Table

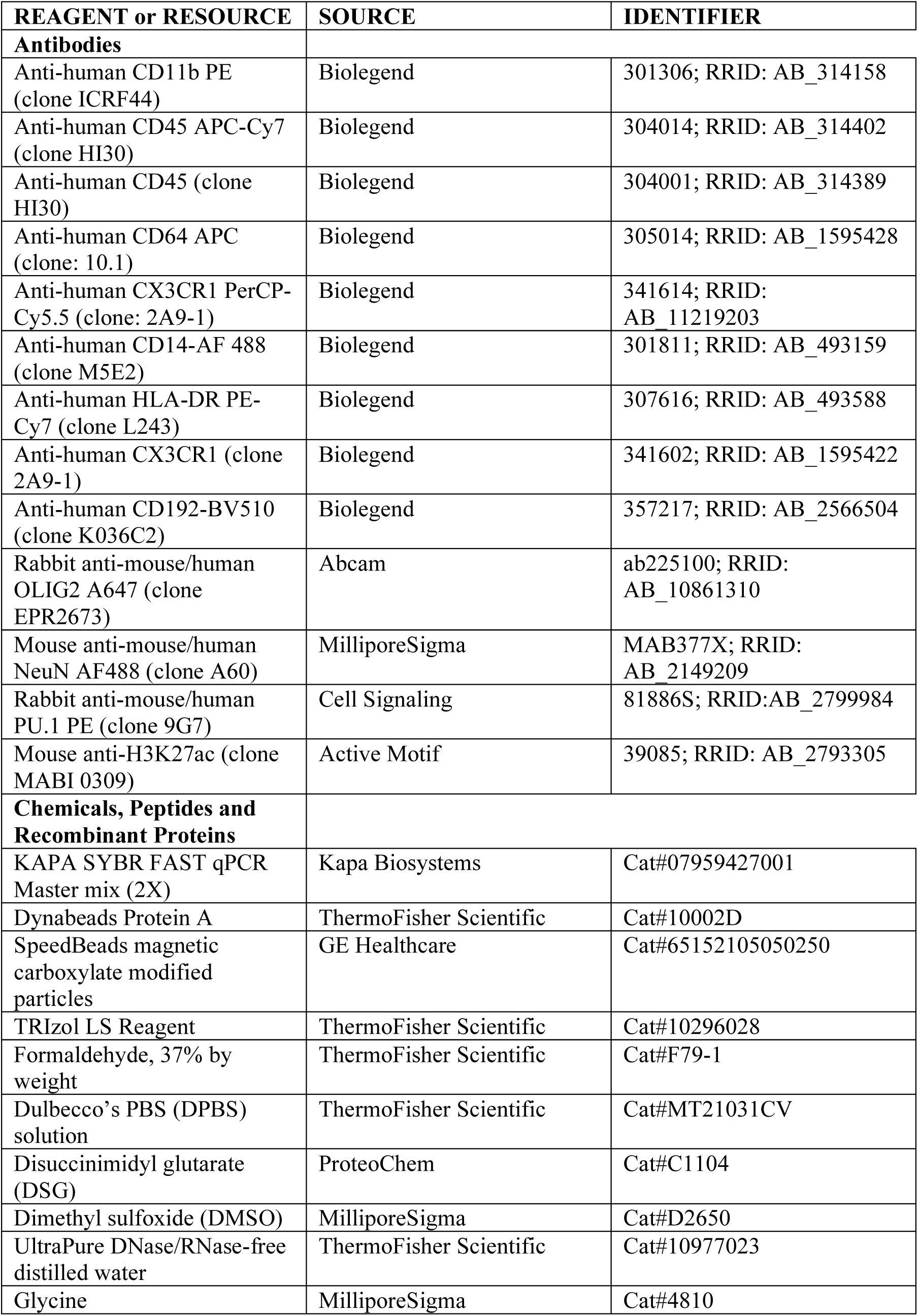
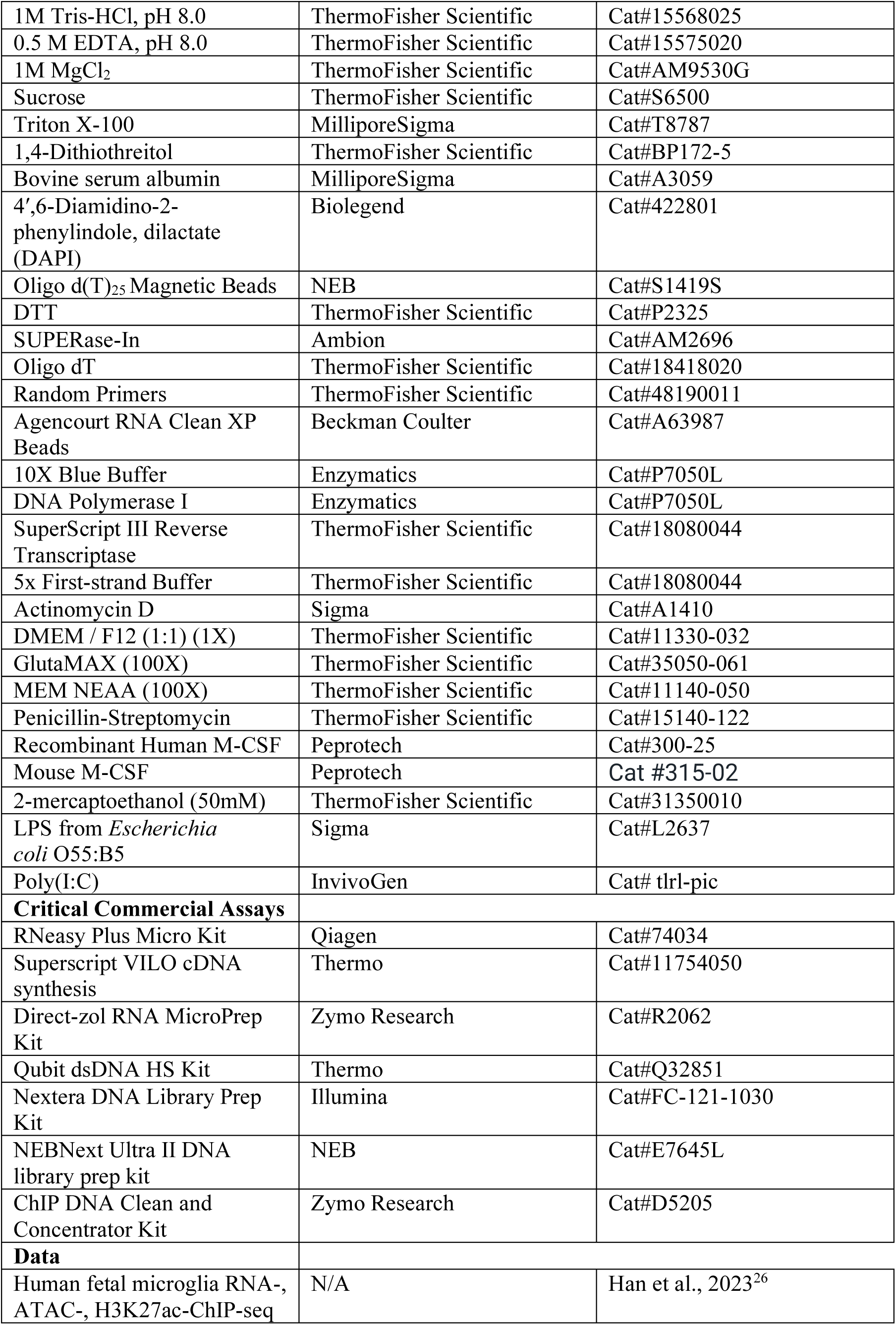
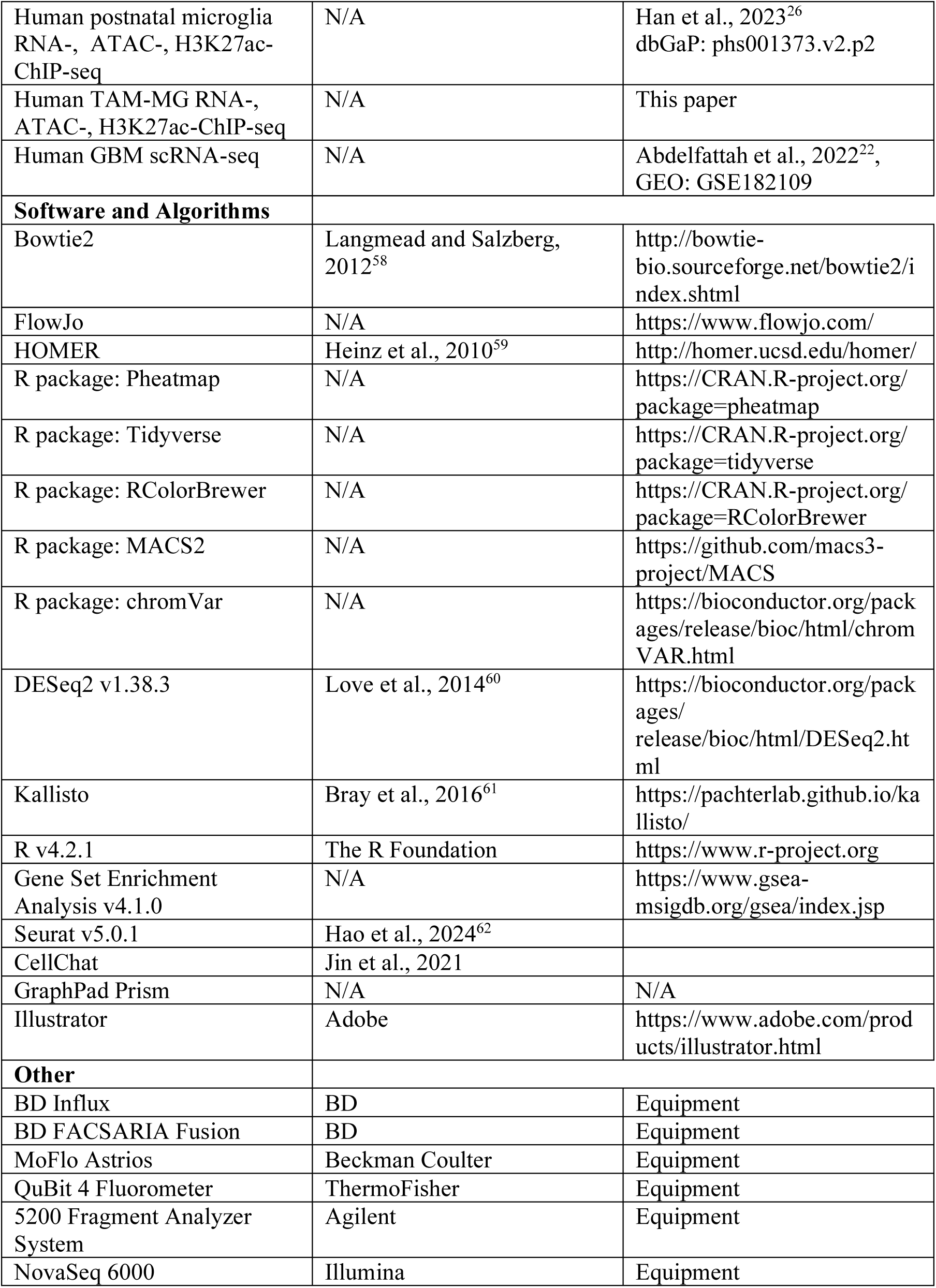

### Resource Availability

#### Lead contact

Further information and requests for resources and reagents should be directed to and will be fulfilled by the lead contact, David. C. Page.

#### Materials availability

This study did not generate new unique reagents.

#### Data and code availability

● Raw RNA-Seq, ATAC-Seq, and ChIP-Seq data has been deposited to dbGaP and processed data has been deposited at github. Both are publicly available as of the date of publication. Accession numbers and DOIs are listed in the key resources table.
● Original code has been deposited at github and is publicly available as of the date of publication. The accession number is listed in the key resources table.
● Any additional information required to reanalyze the data reported in this paper is available from the lead contact upon request.

### Experimental Model and Study Participant Details

#### Human tissue

Isolation of microglia was performed as previously described from brain tissue in excess of that required for diagnosis of pathology. For control microglia samples, all patients were undergoing surgery for epilepsy and epileptogenic focus resections. Surgeries were performed at Rady Children’s Hospital or through University of California (UC) San Diego Health (Jacobs Medical Center or UC San Diego Medical Center Hillcrest). All tumor tissue resections were performed at UC San Diego Hospital. Adult patient consent was obtained for brain tissue and was approved under a protocol by the UC San Diego and Rady Children’s Hospital Institutional Review Board (IRB 160531, IRB 171361). Brain tissue resections were transferred to the laboratory on ice and microglia isolation was performed within three hours after resection. Patient charts were reviewed prior to surgery to confirm pathological diagnosis, medications, demographics, and timing of stereoelectroencephalography. This study was performed in accordance with ethical and legal guidelines of the UC institutional review board. Cell viability and sequencing libraries reported in this study met technical quality control standards and no other criteria were used to exclude samples. We complied with all relevant ethical regulations.

#### Mice

Wild-type C57BL/6J and 129S4/SvJaeJ mice were acquired from the Jackson Laboratory. The mice used in this study were bred and maintained at the Whitehead Institute. All animals were maintained and procedures performed in accordance with the guidelines of the Massachusetts Institute of Technology (MIT) Division of Comparative Medicine, which is overseen by MIT’s Institutional Animal Care and Use Committee (IACUC). The animal care program at MIT/Whitehead Institute is accredited by the Association for Assessment and Accreditation of Laboratory Animal Care, International (AAALAC) and meets or exceeds the standards of AAALAC as detailed in the Guide for the Care and Use of Laboratory Animals. The MIT IACUC approved this research (no. 230-4000-510).

### Method Details

#### Human microglia isolation

Dissection of human brain tissues into 2-3 mm pieces was done manually. Tissue pieces were immersed in homogenization buffer (HBSS, Life Technologies), 1% bovine serum albumin (Sigma-Aldrich, 1 mM EDTA) and mechanically dissociated using a 2 ml polytetrafluoroethylene pestle (Wheaton). Brain homogenate was pelleted, filtered through 40 µm filter, re-suspended in 37% isotonic Percoll (Sigma) and centrifuged at 600xg for 30 min at 16-18°C with minimal acceleration and no deceleration. Percoll enrichment was performed and pelleted cells were collected. Red blood cells were lysed (eBioscience). Remaining cells were washed twice with homogenization buffer and filtered with a 40 µm strainer (BD Falcon). Incubation with Fc-receptor blocking antibody (Human TruStain FcX, BioLegend) in homogenization buffer for 20 minutes on ice was performed. For FACS purification, cells were stained for 30 minutes on ice with the following cell surface marker antibodies at 1:100 dilution (BioLegend): CD11b-PE (301306, clone ICRF44,), CD45-APC/Cy7 (304014, clone HI30), CD64-APC (305014, clone 10.1), CX3CR1-PerCP/Cy5.5 (341614, clone 2A9-1), CD14-AF 488 (301811, clone M5E2), HLA-DR-PE/Cy7 (307616, clone L243), and CD192-BV510 (357217, clone K036C2). Viable cells were first gated using Zombie Violet (Biolegend) or DAPI and added just prior to sorting (1 µg/ml final concentration). A BD Influx (100-µm nozzle, 22 PSI, 2-drop purity mode, sample chilling) or BD FACS AriaFusion (100-µm nozzle, 20 PSI, purity mode, 1-2 drop sort mode, sample chilling) were used to sort microglia defined as live/DAPI^-^/Zombie violet^-^; CD11b^+^; CD45^Low^; CD64^+^; CX3CR1^High^; CD192-BV510^Low^ single cells. FlowJo software (Tree Star) was used to analyze FACS data.

#### Bulk RNA sequencing (RNA-seq)

Microglia post-FACS sorting were stored in TRIzol LS. Phenol-chloroform extraction was used to isolate total RNA from homogenates and stored at -80°C until cDNA libraries were prepared for RNA-seq. We prepared RNA-seq libraries as previously described^25^. mRNAs were incubated with Oligo d(T) Magnetic Beads (New England BioLabs) and fragmented in 2x Superscript III first-strand buffer (ThermoFisher Scientific) with 10mM DTT (ThermoFisher Scientific) at 94°C for 9 minutes. Fragment mRNA was incubated with 0.5 μl of Random primers (3 mg/mL) (ThermoFisher Scientific), 0.5 μl of 50mM Oligo dT primer, (ThermoFisher Scientific), 0.5 μl of SUPERase-In (ThermoFisher Scientific), 1 μl of dNTPs (10 mM) at 50°C for one minute. Then, 1 μl of 10mM DTT, 6 μl of H_2_O+0.02%Tween-20 (Sigma), 0.1 μl Actinomycin D (2 mg/mL), and 0.5 μl of Superscript III (ThermoFisher Scientific) were added to the mixture. Synthesis of cDNA was performed by incubating the resulting mixture in a PCR machine with the following program: 25°C for 10 minutes, 50°C for 50 minutes, and a 4°C hold. RNAClean XP beads (Beckman Coulter) were used to purify the product according to manufacturer’s instructions and eluted with 10 μl of nuclease-free H_2_O. Resulting elution was then incubated with 1.5 μl of Blue Buffer (Enzymatics), 1.1 μl of dUTP mix (10 mM dATP, dCTP, dGTP and 20 mM dUTP), 0.2 mL of RNase H (5 U/mL), 1.2 μl of H_2_O+0.02%Tween-20, and 1 μl of DNA polymerase I (Enzymatics) at 16°C overnight. Purification of DNA was executed using 3 μl of SpeedBeads (ThermoFisher Scientific) resuspended in 28 μl of 20% PEG8000/2.5M NaCl to a final concentration of 13% PEG. Elution of DNA with 40 mL nuclease free H_2_O+0.02%Tween-20 was performed followed by end repair by blunting, A-tailing and adaptor ligation as previously described^54^ using barcoded adapters. PCR amplification of libraries was carried out for 12-15 cycles and a 200-500 bp product size was selected by gel extraction. 51 cycles of sequencing were performed on a HiSeq 4000 (Illumina) or a NextSeq 500 (Illumina).

#### Assay for Transposase-Accessible Chromatin sequencing (ATAC-Seq)

Human microglia (30,000-50,000) were lysed in 50 µl lysis buffer (10 mM Tris-HCl pH 7.5, 10 mM NaCl, 3 mM MgCl_2_, 0.1% IGEPAL, CA-630, in water). Nuclei were centrifuged at 500 rcf for 10 minutes. Pelleted nuclei were resuspended in 50 µl transposase reaction mix (1x Tagment DNA buffer [Illumina], 2.5 µl Tagment DNA enzyme I [Illumina], and incubated at 37°C for 30 min on a heat block. Microglia were directly placed in 50 µl transposase reaction mix for isolations resulting in under 30,000 microglia and incubated for 37°C for 30 min. Zymo ChIP DNA concentrator columns (Zymo Research) were used to purify DNA, followed by elution with 11 µl of elution buffer, and amplification using NEBNext High-Fidelity 2x PCR MasterMix (New England BioLabs) with the Nextera primer Ad1 (1.25 µM) and a unique Ad2.n barcoding primer (1.25 µM) for 8-12 cycles. Size-selection of libraries was performed by gel excision for fragments that were 175-255 bp. Single-end sequencing was performed for 51 cycles on a HiSeq 4000 or NextSeq 500.

#### Chromatin immunoprecipitation-sequencing (ChIP-Seq)

FACS-sorted microglia were centrifugated at 300 rcf and resuspended in 1% PFA. Microglia were rocked for 10 minutes at room temperature. Quenching of PFA was performed using 2.625M glycine at 1:20 volume for 10 minutes at room temperature. Fixed microglia were washed two times and centrifuged at 800-1000 rcf for 5 minutes. Pellets were snap frozen in liquid nitrogen. Snap-frozen microglia pellets containing 250,000 to 500,000 cells were thawed on ice and resuspended using 130 µl of LB3 buffer (10 mM TrisHCl pH 7.5, 100 mM NaCl, 1 mM EDTA, 0.5 mM EGTA, 0.1% Na-Deoxycholate, 0.5% N-Lauroylsarcosine, 1x protease inhibitors). Microglia were transferred to AFA Fiber microtubes (Covaris, MA). Sonication was performed using a Covaris E220 focused-ultrasonicator (Covaris, MA) for 12 cycles of 60 secs (Duty: 5, PIP: 140, Cycles: 200, AMP/Vel/Dwell: 0.0). Post-sonication, samples were transferred to an Eppendorf tube. Triton X-100 was added to the sample for a final concentration of 1%. Supernatant was spun at 21,000 rcf and the pellet discarded. 1% of the total volume was saved as DNA input control and stored at -20°C until library preparation. 25 µl of Protein A DynaBeads (ThermoFisher Scientific) and 1 µl of H3K27ac antibody (Active Motif) were added to the supernatant for the immunoprecipitation. Samples were rotated at 4°C overnight. Dynabeads were washed 3 times with Wash Buffer 1 (20 mM Tris-HCl pH 7.4, 150 mM NaCl, 2 mM EDTA, 0.1% SDS, 1% Triton X-100), three times with Wash Buffer 3 (10 mM Tris-HCl pH, 250 mM LiCl, 1 mM EDTA, 1% Triton X100, 0.7% Na-Deoxycholate), three times with TET (10 mM Tris-HCl pH 8, 1 mM EDTA, 0.2% Tween20), once with TE-NaCl (10 mM Tris-HCl pH 8, 1 mM EDTA, 50 mM NaCl) and resuspended in 25 µl TT (10 mM Tris-HCl pH 8, 0.05% Tween20). Input samples were adjusted to 25 µl with TT. NEBNext Ultra II DNA Library Prep kit (New England BioLabs E7645) was used to prepare sample and input libraries according to manufacturer’s instructions. Samples and inputs were de-crosslinked (RNase A, Proteinase K, and 4.5 µl of 5M NaCl) and incubated overnight at 65°C. PCR-amplification of libraries was performed using NEBNext High Fidelity 2X PCR MasterMix (New England BioLabs) for 14 cycles. Size selection of libraries was performed by gel excision of fragments that were 225 to 500 bp. Single-end sequencing of libraries for 51 cycles on a HiSeq 4000 or NextSeq 500 was performed.

#### Mouse microglia isolation

Mouse microglia were isolated as previously described using gentle mechanical dissociation and a 37% isopercoll cushion^25,30^. Microglia were then incubated in staining buffer on ice with anti-CD16/32 blocking antibody (BioLegend 101319, 1:500) for 15 min. Anti-mouse anti-CD11b-APC (BioLegend 101212, 1:100), anti-CD45-Alexa488 (BioLegend 103122, 1:100), and anti-CX3CR1-PE (BioLegend 149006, 1:100) were then added to microglia for 25 min or overnight on ice.

#### Microglia in vitro culture and stimulation

Post FACS, microglia were resuspended in media containing 20ng/ml M-CSF and plated at 50,000 cells per well in a 96-well plate. Microglia were incubated at 37°C for at least two days prior to stimulation. In vitro-cultured microglia were treated for 4 hours with 100 ng/ml LPS or 10 ug/ml poly (I:C). Cells were washed and collected for RNA isolation and quantitative RT-PCR or RNA-seq.

#### Quantitative RT-PCR

500 ng RNA was added to Superscript VILO cDNA synthesis reaction (ThermoFisher). cDNA was diluted 1:3 and 3ul was added to the qRT-PCR reaction using 2X Power SYBR green qPCR master mix (ThermoFisher) for total volume of 10 uL. Endogenous control genes *Actb* and *Gapdh* were used. Three technical replicates were quantified per sample.

### Data analysis

#### Bulk RNA-seq

All human analyses were performed using human genome build hg38, and a custom version of the comprehensive GENCODE v24 transcriptome annotation described in San Roman et al., 2023. Reads were pseudoaligned to the transcriptome annotation, and expression levels of each transcript were estimated using kallisto software v0.42.5. Resulting count data were imported into R with the Tximport package v1.14.0 for normalization using DESeq2 v1.26.0. Downstream analysis used only protein-coding genes (as annotated in ensembl v104) with exceptions described in San Roman et al., 2023. All mouse analyses were performed using mouse genome build mm10, and GENCODE vM15 transcriptome annotation. Reads were pseudoaligned and transcript counts estimated and normalized as described for human samples. Differentially expressed genes (DEGs) between TAM-MGs and control microglia, and between XX and XY TAM-MGs for grade II-III and GBM, were identified using DESeq2. For TAM-MG vs. control microglia, we used a cutoff of log_2_FC>0.58, adjusted-p<0.05. For sex-biased genes, we used a cutoff of log_2_FC>0.58, p<0.05.

#### Gene set enrichment analysis

Gene set enrichment analysis was conducted using GSEA version 4.1.0 software and the 50 Hallmark pathways were downloaded from the Molecular Signatures Database. Analysis was restricted to autosomal protein-coding and lincRNA genes, which were ranked by each gene’s t-statistic from the DESeq2 models for TAM-MG vs control or XX vs XY comparisons. Results were considered statistically significant if FDR<0.05.

#### ATAC-seq and ChIP-seq analysis

Peaks were called using HOMER’s findPeaks command with the following parameters: ‘‘-style factor - size 200 -minDist 200’’ for ATAC-seq experiments and ‘‘-style histone -size 500 -minDist 1000 -region’’ for ChIP-seq experiments. Peaks were merged with HOMER’s mergePeaks and annotated using HOMER’s annotatePeaks.pl using all tag directories. For ChIP-seq experiments, peaks were annotated around ATAC-seq peaks with the parameter ‘‘-size -500,500 -pc’’. Subsequently, DESeq292 was used to identify the differentially chromatin accessible distal sites (1000bp away from known TSS) or proximal sites (<500bp away from known transcript) with p-adj <0.05 and fold change >2.

#### Motif analysis

De novo motif analysis was performed using HOMER’s findMotifsGenome.pl with either all peaks or random genome sequences as background peaks. Motif enrichment scoring was performed using binomial distribution under HOMER’s framework.

#### Single-cell RNA sequencing

Single cell RNA-seq FASTQ files were aligned using CellRanger (v7.1.0) and CellRanger’s pre-built reference genome for human (hg38). Cells were used that met the quality control metrics of percent mitochondrial gene expression < 5% and number of expressed genes per cell (nFeature) > 500 and < 2500. Each GBM sample was merged into one Seurat (v5.0.1) object, and integrated using Harmony. Seurat clustering was performed and cell types were annotated based on known marker genes. We identified genes differentially expressed in XX and XY samples for each of the four cell types analyzed: TAM-MGs, TAM-BMDMs, T-cells, and tumor cells. The minimum detection rate for a given differentially expressed gene across cell populations = 0.2. We applied the established method CellChat to predict interactions between the four cell types based on sex-biased enrichment of the manually curated CellChat ligand-receptor interaction database.

### Statistical analyses

Various statistical tests were used to calculate p-values as indicated in the Methods Details, figure legends, or text. To calculate statistics and generate plots, we used R software, version 4.2.1. Gene expression differences were calculated with DESeq2 with Benjamini-Hochberg multiple testing correction. We considered results statistically significant when p<0.05 or, when using multiple hypothesis correction, adjusted-p<0.05 or FDR<0.05.

### Data Visualization

PCA and heatmaps were generated in R and other plots were made with ggplots2 in R with colors reflecting the scores/expression values, including z-scores, as noted in each figure. Browser images were generated from the UCSC Genome Browser.

## Acknowledgements

We thank members of the Page laboratory for critical discussions on the project narrative and experimental design. We thank Jorge Adarme, Susan Tocio, Laura Brown, and Alexis Drake for laboratory support. We thank Jennifer Hughes, Alisa White, Winston Bellott, Rebecca Harris, and Maya Talukdar for manuscript editing. We thank the Whitehead Institute FACS Core facility and Genome Technology Core facility for cell sorting, library preparation and sequencing. We thank Caitlin Rausch for her contributions to figure illustrations. We thank the individuals who contributed tissue samples for their participation.

Funding: Jane Coffin Childs Fund (M.E.T.), Koch Institute Frontier Research Fund (M.E.T.), Howard Hughes Medical Institute (D.C.P.), and Simons Foundation Autism Research Initiative award 809293 (D.C.P., C.K.G., N.G.C). Philanthropic support from The Brit Jepson d’Arbeloff Center on Women’s Health, Arthur W. and Carol Tobin Brill, Matthew Brill, Charles Ellis, Carla Knobloch, The Brett Barakett Foundation, Howard P. Colhoun Family Foundation, and the Seedlings Foundation. C.Z.H. is supported by the Cancer Research Institute Irvington Postdoctoral Fellowship Program and NIH K99 MH129983. N.G.C. and C.K.G. are supported by NIH R01 NS096170 and the Chan Zuckerberg Initiative. C.K.G. is supported by the JPB foundation grant KR29574. N.G.C. is supported by NIH grants K08 NS109200, R01 NS124637, NS126452, the Doris Duke foundation and the Hartwell Foundation. C.K.G. are supported by the Cure Alzheimer’s Fund, and Alzheimer’s Association ADSF-21-829655-C. The Flow Cytometry Core Facility of the Salk Institute is partly supported by NIH-NCI CCSG: P30 014195 and SIG S10-OD023689. This publication includes data generated at the UC San Diego IGM Genomics Center Illumina NovaSeq 6000 purchased with NIH SIG grant (S10 OD026929). A.N. is supported by the UK Dementia Research Institute [award number UKDRI-5016] through UK DRI Ltd, principally funded by the Medical Research Council. This study was partly funded through a collaboration with Celgene / BMS.

## Author contributions

Conceptualization: M.E.T., C.Z.H, and D.C.P. Data curation: M.E.T. and C.Z.H; Patient identification, consent, and tissue acquisition: C.F., C.O., S.P., J.B., L.E., K.M., M.G., M.S.S., H.S.U., P.S.J., M.L.L., D.D.G., S.B.H., J.C., D.B., A.K., N.G.C., C.C.C.; Sorting and sequencing: C.Z.H., C.D.B., A.N.; Formal analysis: M.E.T. and C.Z.H; Funding acquisition: M.E.T., C.Z.H, C.K.G., and D.C.P.; Writing—original draft preparation: M.E.T., C.Z.H., and D.C.P.; Writing—review and editing: M.E.T., C.Z.H., C.K.G., and D.C.P.

## Inclusion and diversity

We support inclusive, diverse, and equitable conduct of research.

## References

1. Ostrom, Q.T., Price, M., Neff, C., Cioffi, G., Waite, K.A., Kruchko, C., and Barnholtz-Sloan, Jill S. (2022). CBTRUS Statistical Report: Primary Brain and Other Central Nervous System Tumors Diagnosed in the United States in 2015–2019. Neuro-Oncology 24, v1–v95. 10.1093/neuonc/noac202.

2. Khabibov, M., Garifullin, A., Boumber, Y., Khaddour, K., Fernandez, M., Khamitov, F., Khalikova, L., Kuznetsova, N., Kit, O., and Kharin, L. (2022). Signaling pathways and therapeutic approaches in glioblastoma multiforme (Review). Int J Oncol 60. 10.3892/ijo.2022.5359.

3. Poon, M.T.C., Sudlow, C.L.M., Figueroa, J.D., and Brennan, P.M. (2020). Longer-term (≥ 2 years) survival in patients with glioblastoma in population-based studies pre- and post-2005: a systematic review and meta-analysis. Scientific Reports 10, 11622. 10.1038/s41598-020-68011-4.

4. Liu, Y., Zhou, F., Ali, H., Lathia, J.D., and Chen, P. (2024). Immunotherapy for glioblastoma: current state, challenges, and future perspectives. Cellular & Molecular Immunology. 10.1038/s41423-024-01226-x.

5. Choi, B.D., Gerstner, E.R., Frigault, M.J., Leick, M.B., Mount, C.W., Balaj, L., Nikiforow, S., Carter, B.S., Curry, W.T., Gallagher, K., and Maus, M.V. (2024). Intraventricular CARv3-TEAM-E T Cells in Recurrent Glioblastoma. New England Journal of Medicine. 10.1056/NEJMoa2314390.

6. Himes, B.T., Geiger, P.A., Ayasoufi, K., Bhargav, A.G., Brown, D.A., and Parney, I.F. (2021). Immunosuppression in Glioblastoma: Current Understanding and Therapeutic Implications. Front Oncol 11, 770561. 10.3389/fonc.2021.770561.

7. O’Donnell, J.S., Teng, M.W.L., and Smyth, M.J. (2019). Cancer immunoediting and resistance to T cell-based immunotherapy. Nature Reviews Clinical Oncology 16, 151–167. 10.1038/s41571-018-0142-8.

8. Ochocka, N., Segit, P., Wojnicki, K., Cyranowski, S., Swatler, J., Jacek, K., Grajkowska, W., and Kaminska, B. (2023). Specialized functions and sexual dimorphism explain the functional diversity of the myeloid populations during glioma progression. Cell Rep 42, 111971. 10.1016/j.celrep.2022.111971.

9. Dubinski, D., Wölfer, J., Hasselblatt, M., Schneider-Hohendorf, T., Bogdahn, U., Stummer, W., Wiendl, H., and Grauer, O.M. (2015). CD4+ T effector memory cell dysfunction is associated with the accumulation of granulocytic myeloid-derived suppressor cells in glioblastoma patients. Neuro-Oncology 18, 807–818. 10.1093/neuonc/nov280.

10. Wang, L., Jung, J., Babikir, H., Shamardani, K., Jain, S., Feng, X., Gupta, N., Rosi, S., Chang, S., Raleigh, D., et al. (2022). A single-cell atlas of glioblastoma evolution under therapy reveals cell-intrinsic and cell-extrinsic therapeutic targets. Nature Cancer 3, 1534–1552. 10.1038/s43018-022-00475-x.

11. Ostrom, Q.T., Rubin, J.B., Lathia, J.D., Berens, M.E., and Barnholtz-Sloan, J.S. (2018). Females have the survival advantage in glioblastoma. Neuro Oncol 20, 576–577. 10.1093/neuonc/noy002.

12. Klein, S.L., and Flanagan, K.L. (2016). Sex differences in immune responses. Nature Reviews Immunology 16, 626–638. 10.1038/nri.2016.90.

13. Sun, T., Warrington, N.M., Luo, J., Brooks, M.D., Dahiya, S., Snyder, S.C., Sengupta, R., and Rubin, J.B. (2014). Sexually dimorphic RB inactivation underlies mesenchymal glioblastoma prevalence in males. J Clin Invest 124, 4123–4133. 10.1172/jci71048.

14. Wang, G., Zhong, K., Wang, Z., Zhang, Z., Tang, X., Tong, A., and Zhou, L. (2022). Tumor-associated microglia and macrophages in glioblastoma: From basic insights to therapeutic opportunities. Front Immunol 13, 964898. 10.3389/fimmu.2022.964898.

15. Paolicelli, R.C., Sierra, A., Stevens, B., Tremblay, M.E., Aguzzi, A., Ajami, B., Amit, I., Audinat, E., Bechmann, I., Bennett, M., et al. (2022). Microglia states and nomenclature: A field at its crossroads. Neuron 110, 3458–3483. 10.1016/j.neuron.2022.10.020.

16. Holtman, I.R., Skola, D., and Glass, C.K. (2017). Transcriptional control of microglia phenotypes in health and disease. J Clin Invest 127, 3220–3229. 10.1172/jci90604.

17. Greenwald, A.C., Darnell, N.G., Hoefflin, R., Simkin, D., Mount, C.W., Gonzalez Castro, L.N., Harnik, Y., Dumont, S., Hirsch, D., Nomura, M., et al. (2024). Integrative spatial analysis reveals a multi-layered organization of glioblastoma. Cell. 10.1016/j.cell.2024.03.029.

18. Sørensen, M.D., Dahlrot, R.H., Boldt, H.B., Hansen, S., and Kristensen, B.W. (2018). Tumour-associated microglia/macrophages predict poor prognosis in high-grade gliomas and correlate with an aggressive tumour subtype. Neuropathol Appl Neurobiol 44, 185–206. 10.1111/nan.12428.

19. Sørensen, M.D., and Kristensen, B.W. (2022). Tumour-associated CD204(+) microglia/macrophages accumulate in perivascular and perinecrotic niches and correlate with an interleukin-6-enriched inflammatory profile in glioblastoma. Neuropathol Appl Neurobiol 48, e12772. 10.1111/nan.12772.

20. Liu, H., Sun, Y., Zhang, Q., Jin, W., Gordon, R.E., Zhang, Y., Wang, J., Sun, C., Wang, Z.J., Qi, X., et al. (2021). Pro-inflammatory and proliferative microglia drive progression of glioblastoma. Cell Rep 36, 109718. 10.1016/j.celrep.2021.109718.

21. Turaga, S.M., Silver, D.J., Bayik, D., Paouri, E., Peng, S., Lauko, A., Alban, T.J., Borjini, N., Stanko, S., Naik, U.P., et al. (2020). JAM-A functions as a female microglial tumor suppressor in glioblastoma. Neuro Oncol 22, 1591–1601. 10.1093/neuonc/noaa148.

22. Abdelfattah, N., Kumar, P., Wang, C., Leu, J.S., Flynn, W.F., Gao, R., Baskin, D.S., Pichumani, K., Ijare, O.B., Wood, S.L., et al. (2022). Single-cell analysis of human glioma and immune cells identifies S100A4 as an immunotherapy target. Nat Commun 13, 767. 10.1038/s41467-022-28372-y.

23. Louis, D.N., Perry, A., Wesseling, P., Brat, D.J., Cree, I.A., Figarella-Branger, D., Hawkins, C., Ng, H.K., Pfister, S.M., Reifenberger, G., et al. (2021). The 2021 WHO Classification of Tumors of the Central Nervous System: a summary. Neuro Oncol 23, 1231–1251. 10.1093/neuonc/noab106.

24. Komohara, Y., Ohnishi, K., Kuratsu, J., and Takeya, M. (2008). Possible involvement of the M2 anti-inflammatory macrophage phenotype in growth of human gliomas. J Pathol 216, 15–24. 10.1002/path.2370.

25. Gosselin, D., Skola, D., Coufal, N.G., Holtman, I.R., Schlachetzki, J.C.M., Sajti, E., Jaeger, B.N., O’Connor, C., Fitzpatrick, C., Pasillas, M.P., et al. (2017). An environment-dependent transcriptional network specifies human microglia identity. Science 356. 10.1126/science.aal3222.

26. Han, C.Z., Li, R.Z., Hansen, E., Trescott, S., Fixsen, B.R., Nguyen, C.T., Mora, C.M., Spann, N.J., Bennett, H.R., Poirion, O., et al. (2023). Human microglia maturation is underpinned by specific gene regulatory networks. Immunity 56, 2152–2171.e2113. 10.1016/j.immuni.2023.07.016.

27. Liberzon, A., Birger, C., Thorvaldsdóttir, H., Ghandi, M., Mesirov, J.P., and Tamayo, P. (2015). The Molecular Signatures Database (MSigDB) hallmark gene set collection. Cell Syst 1, 417–425. 10.1016/j.cels.2015.12.004.

28. Ha, E.T., Antonios, J.P., Soto, H., Prins, R.M., Yang, I., Kasahara, N., Liau, L.M., and Kruse, C.A. (2014). Chronic inflammation drives glioma growth: cellular and molecular factors responsible for an immunosuppressive microenvironment. Neuroimmunology and Neuroinflammation 1, 66–76. 10.4103/2347-8659.139717.

29. Gudgeon, J., Marín-Rubio, J.L., and Trost, M. (2022). The role of macrophage scavenger receptor 1 (MSR1) in inflammatory disorders and cancer. Front Immunol 13, 1012002. 10.3389/fimmu.2022.1012002.

30. Fixsen, B.R., Han, C.Z., Zhou, Y., Spann, N.J., Saisan, P., Shen, Z., Balak, C., Sakai, M., Cobo, I., Holtman, I.R., et al. (2023). SALL1 enforces microglia-specific DNA binding and function of SMADs to establish microglia identity. Nature Immunology 24, 1188–1199. 10.1038/s41590-023-01528-8.

31. Keren-Shaul, H., Spinrad, A., Weiner, A., Matcovitch-Natan, O., Dvir-Szternfeld, R., Ulland, T.K., David, E., Baruch, K., Lara-Astaiso, D., Toth, B., et al. (2017). A Unique Microglia Type Associated with Restricting Development of Alzheimer’s Disease. Cell 169, 1276–1290.e1217. 10.1016/j.cell.2017.05.018.

32. Agostini, F., Agostinis, R., Medina, D.L., Bisaglia, M., Greggio, E., and Plotegher, N. (2022). The Regulation of MiTF/TFE Transcription Factors Across Model Organisms: from Brain Physiology to Implication for Neurodegeneration. Mol Neurobiol 59, 5000–5023. 10.1007/s12035-022-02895-3.

33. Li, J., and Stanger, B.Z. (2020). How Tumor Cell Dedifferentiation Drives Immune Evasion and Resistance to Immunotherapy. Cancer Res 80, 4037–4041. 10.1158/0008-5472.Can-20-1420.

34. Rock, R.B., Hu, S., Deshpande, A., Munir, S., May, B.J., Baker, C.A., Peterson, P.K., and Kapur, V. (2005). Transcriptional response of human microglial cells to interferon-gamma. Genes Immun 6, 712–719. 10.1038/sj.gene.6364246.

35. Chearwae, W., and Bright, J.J. (2008). PPARgamma agonists inhibit growth and expansion of CD133+ brain tumour stem cells. Br J Cancer 99, 2044–2053. 10.1038/sj.bjc.6604786.

36. Straus, D.S., and Glass, C.K. (2007). Anti-inflammatory actions of PPAR ligands: new insights on cellular and molecular mechanisms. Trends Immunol 28, 551–558. 10.1016/j.it.2007.09.003.

37. Tao, T., Wang, Y., Luo, H., Yao, L., Wang, L., Wang, J., Yan, W., Zhang, J., Wang, H., Shi, Y., et al. (2013). Involvement of FOS-mediated miR-181b/miR-21 signalling in the progression of malignant gliomas. European Journal of Cancer 49, 3055–3063. 10.1016/j.ejca.2013.05.010.

38. Jin, S., Guerrero-Juarez, C.F., Zhang, L., Chang, I., Ramos, R., Kuan, C.-H., Myung, P., Plikus, M.V., and Nie, Q. (2021). Inference and analysis of cell-cell communication using CellChat. Nature Communications 12, 1088. 10.1038/s41467-021-21246-9.

39. Chen, D., Varanasi, S.K., Hara, T., Traina, K., Sun, M., McDonald, B., Farsakoglu, Y., Clanton, J., Xu, S., Garcia-Rivera, L., et al. (2023). CTLA-4 blockade induces a microglia-Th1 cell partnership that stimulates microglia phagocytosis and anti-tumor function in glioblastoma. Immunity 56, 2086–2104.e2088. 10.1016/j.immuni.2023.07.015.

40. San Roman, A.K., Godfrey, A.K., Skaletsky, H., Bellott, D.W., Groff, A.F., Harris, H.L., Blanton, L.V., Hughes, J.F., Brown, L., Phou, S., et al. (2023). The human inactive X chromosome modulates expression of the active X chromosome. Cell Genom 3, 100259. 10.1016/j.xgen.2023.100259.

41. Blanton, L.V., San Roman, A.K., Wood, G., Buscetta, A., Banks, N., Skaletsky, H., Godfrey, A.K., Pham, T.T., Hughes, J.F., Brown, L.G., et al. (2024). Stable and robust Xi and Y transcriptomes drive cell-type-specific autosomal and Xa responses in vivo and in vitro in four human cell types. Cell Genomics, 100628. 10.1016/j.xgen.2024.100628.

42. Bellott, D.W., Hughes, J.F., Skaletsky, H., Brown, L.G., Pyntikova, T., Cho, T.J., Koutseva, N., Zaghlul, S., Graves, T., Rock, S., et al. (2014). Mammalian Y chromosomes retain widely expressed dosage-sensitive regulators. Nature 508, 494–499. 10.1038/nature13206.

43. Saikruang, W., Ang Yan Ping, L., Abe, H., Kasumba, D.M., Kato, H., and Fujita, T. (2022). The RNA helicase DDX3 promotes IFNB transcription via enhancing IRF-3/p300 holocomplex binding to the IFNB promoter. Scientific Reports 12, 3967. 10.1038/s41598-022-07876-z.

44. Kienes, I., Bauer, S., Gottschild, C., Mirza, N., Pfannstiel, J., Schröder, M., and Kufer, T.A. (2021). DDX3X Links NLRP11 to the Regulation of Type I Interferon Responses and NLRP3 Inflammasome Activation. Front Immunol 12, 653883. 10.3389/fimmu.2021.653883.

45. Shen, H., Yanas, A., Owens, M.C., Zhang, C., Fritsch, C., Fare, C.M., Copley, K.E., Shorter, J., Goldman, Y.E., and Liu, K.F. (2022). Sexually dimorphic RNA helicases DDX3X and DDX3Y differentially regulate RNA metabolism through phase separation. Mol Cell 82, 2588–2603.e2589. 10.1016/j.molcel.2022.04.022.

46. Rengarajan, S., Derks, J., Bellott, D.W., Slavov, N., and Page, D.C. (2025). Post-transcriptional cross- and auto-regulation buffer expression of the human RNA helicases DDX3X and DDX3Y. Genome Res 35, 20–30. 10.1101/gr.279707.124.

47. Bol, G.M., Vesuna, F., Xie, M., Zeng, J., Aziz, K., Gandhi, N., Levine, A., Irving, A., Korz, D., Tantravedi, S., et al. (2015). Targeting DDX3 with a small molecule inhibitor for lung cancer therapy. EMBO Mol Med 7, 648–669. 10.15252/emmm.201404368.

48. Özbay Kurt, F.G., Cicortas, B.A., Balzasch, B.M., De la Torre, C., Ast, V., Tavukcuoglu, E., Ak, C., Wohlfeil, S.A., Cerwenka, A., Utikal, J., and Umansky, V. (2024). S100A9 and HMGB1 orchestrate MDSC-mediated immunosuppression in melanoma through TLR4 signaling. J Immunother Cancer 12. 10.1136/jitc-2024-009552.

49. Ferrarese, R., Joseph, K., Andrieux, G., Haase, I.V., Zanon, F., Kling, E., Izzo, A., Corrales, E., Schwabenland, M., Prinz, M., et al. (2024). ZBTB18 regulates cytokine expression and affects microglia/macrophage recruitment and commitment in glioblastoma. Communications Biology 7, 1472. 10.1038/s42003-024-07144-y.

50. Samir, P., Kesavardhana, S., Patmore, D.M., Gingras, S., Malireddi, R.K.S., Karki, R., Guy, C.S., Briard, B., Place, D.E., Bhattacharya, A., et al. (2019). DDX3X acts as a live-or-die checkpoint in stressed cells by regulating NLRP3 inflammasome. Nature 573, 590–594. 10.1038/s41586-019-1551-2.

51. Lakshmikanth, T., Consiglio, C., Sardh, F., Forlin, R., Wang, J., Tan, Z., Barcenilla, H., Rodriguez, L., Sugrue, J., Noori, P., et al. (2024). Immune system adaptation during gender-affirming testosterone treatment. Nature 633, 155–164. 10.1038/s41586-024-07789-z.

52. Lee, J., Kay, K., Troike, K., Ahluwalia, M.S., and Lathia, J.D. (2022). Sex Differences in Glioblastoma Immunotherapy Response. NeuroMolecular Medicine 24, 50–55. 10.1007/s12017-021-08659-x.

53. Lee, J., Nicosia, M., Hong, E.S., Silver, D.J., Li, C., Bayik, D., Watson, D.C., Lauko, A., Kay, K.E., Wang, S.Z., et al. (2023). Sex-Biased T-cell Exhaustion Drives Differential Immune Responses in Glioblastoma. Cancer Discovery 13, 2090–2105. 10.1158/2159-8290.Cd-22-0869.

54. Asija, S., Chatterjee, A., Goda, J.S., Yadav, S., Chekuri, G., and Purwar, R. (2023). Oncolytic immunovirotherapy for high-grade gliomas: A novel and an evolving therapeutic option. Front Immunol 14, 1118246. 10.3389/fimmu.2023.1118246.

55. Maenner, M.J., Shaw, K.A., Bakian, A.V., Bilder, D.A., Durkin, M.S., Esler, A., Furnier, S.M., Hallas, L., Hall-Lande, J., Hudson, A., et al. (2021). Prevalence and Characteristics of Autism Spectrum Disorder Among Children Aged 8 Years - Autism and Developmental Disabilities Monitoring Network, 11 Sites, United States, 2018. MMWR Surveill Summ 70, 1–16. 10.15585/mmwr.ss7011a1.

56. Coales, I., Tsartsalis, S., Fancy, N., Weinert, M., Clode, D., Owen, D., and Matthews, P.M. (2022). Alzheimer’s disease-related transcriptional sex differences in myeloid cells. Journal of Neuroinflammation 19, 247. 10.1186/s12974-022-02604-w.

57. Carling, G.K., Fan, L., Foxe, N.R., Norman, K., Wong, M.Y., Zhu, D., Corona, C., Razzoli, A., Yu, F., Yarahmady, A., et al. (2024). Alzheimer&#x2019;s disease-linked risk alleles elevate microglial cGAS-associated senescence and neurodegeneration in a tauopathy model. Neuron 112, 3877–3896.e3878. 10.1016/j.neuron.2024.09.006.

58. Langmead, B., and Salzberg, S.L. (2012). Fast gapped-read alignment with Bowtie 2. Nature Methods 9, 357–359. 10.1038/nmeth.1923.

59. Heinz, S., Benner, C., Spann, N., Bertolino, E., Lin, Y.C., Laslo, P., Cheng, J.X., Murre, C., Singh, H., and Glass, C.K. (2010). Simple combinations of lineage-determining transcription factors prime cis-regulatory elements required for macrophage and B cell identities. Mol Cell 38, 576–589. 10.1016/j.molcel.2010.05.004.

60. Love, M.I., Huber, W., and Anders, S. (2014). Moderated estimation of fold change and dispersion for RNA-seq data with DESeq2. Genome Biology 15, 550. 10.1186/s13059-014-0550-8.

61. Bray, N.L., Pimentel, H., Melsted, P., and Pachter, L. (2016). Near-optimal probabilistic RNA-seq quantification. Nature Biotechnology 34, 525–527. 10.1038/nbt.3519.

62. Hao, Y., Stuart, T., Kowalski, M.H., Choudhary, S., Hoffman, P., Hartman, A., Srivastava, A., Molla, G., Madad, S., Fernandez-Granda, C., and Satija, R. (2024). Dictionary learning for integrative, multimodal and scalable single-cell analysis. Nature Biotechnology 42, 293–304. 10.1038/s41587-023-01767-y.

