## Supplemental Figures for "The inactive X chromosome drives sex differences in microglial inflammatory activity in human glioblastoma"

**Supplementary Information:**

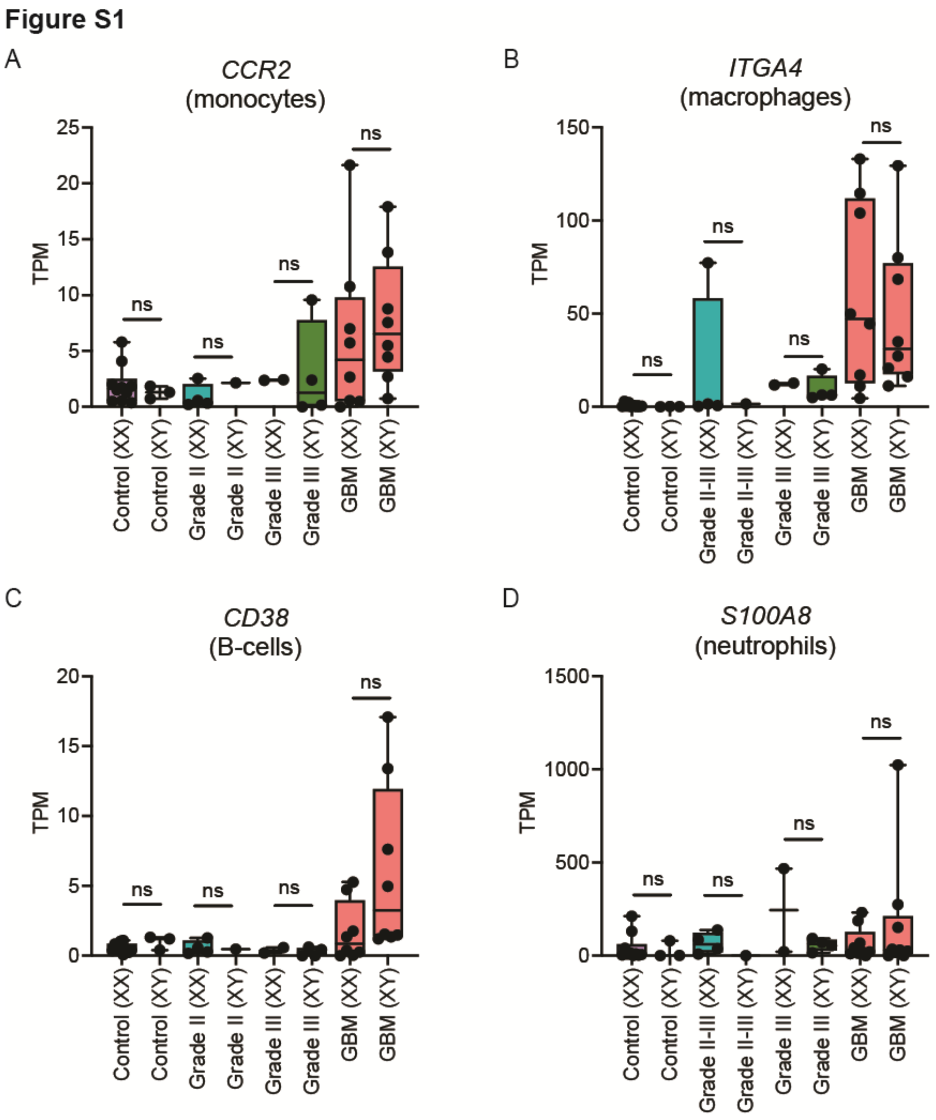


**Supplemental Figure 1.**

A. Expression of monocyte marker *CCR2* across control microglia and TAM-MGs, XX and XY.

B. Expression of blood-derived macrophage marker *ITGA4/C49* across control microglia and TAM-MGs, XX and XY.

C. Expression of B-cell marker *CD38* across control microglia and TAM-MGs, XX and XY.

D. Expression of neutrophil marker *S100A8* across control microglia and TAM-MGs, XX and XY.

### Figure S2

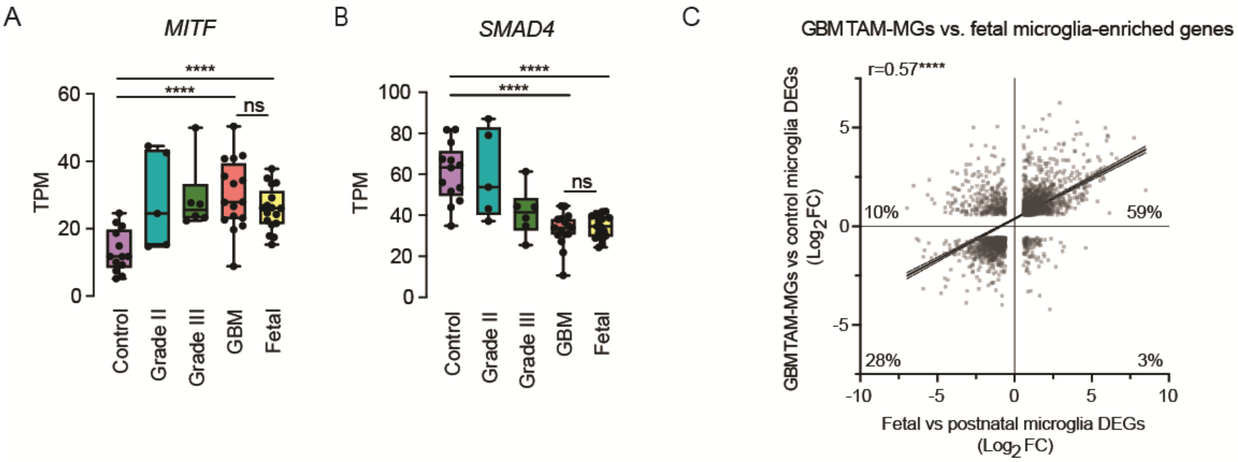


**Supplemental Figure 2.**

A. Expression of GBM TAM-MG and fetal microglia-enriched transcription factor *MITF*.

B. Expression of control microglia and postnatal microglia-enriched transcription factor *SMAD4*.

C. Comparison of DEGs common to GBM TAM-MGs vs control microglia and fetal microglia vs postnatal microglia. DEG = log2FC > 0.58 and padj < 0.05.

**Figure S3**

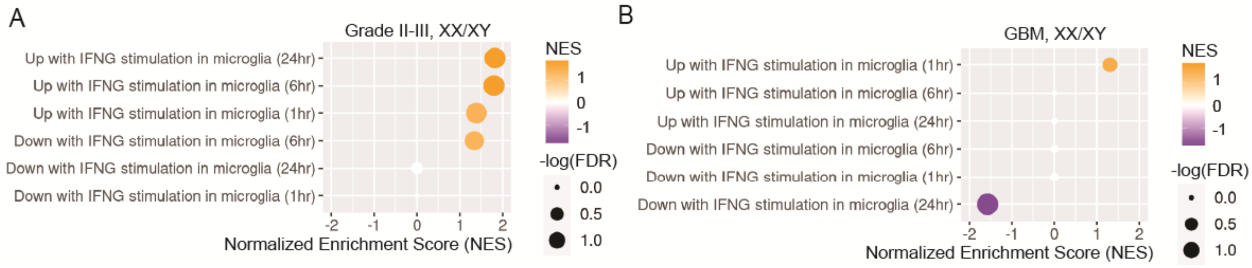


**Supplemental Figure 3.**

A. GSEA of sex-biased genes in grade II-III TAM-MGs belonging to IFN gamma-stimulated gene sets in microglia. NES = normalized enrichment score. FDR = false discovery rate. Pathways with -log (FDR) > 1.3 shown in color.

B. GSEA of sex-biased genes in GBM TAM-MGs belonging to IFN gamma-stimulated gene sets in microglia. NES = normalized enrichment score. FDR = false discovery rate. Pathways with -log (FDR) > 1.3 shown in color.

**Figure S4**

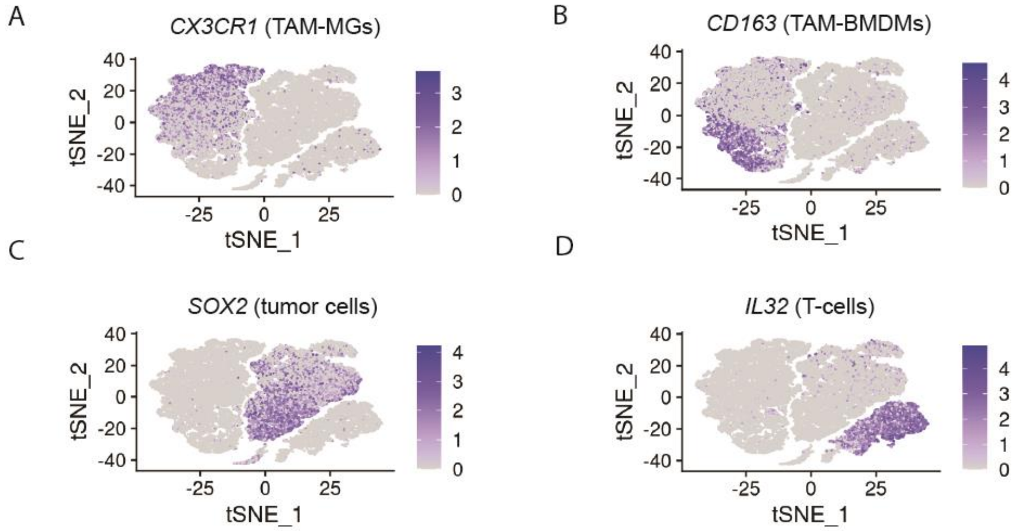


**Supplemental Figure 4.**

A. Expression of TAM-MG marker gene *CX3CR1*.

B. Expression of TAM-BMDM marker gene *CD163*.

C. Expression of tumor cell gene *SOX2*.

D. Expression of T-cell gene *IL32*.

### Figure S5

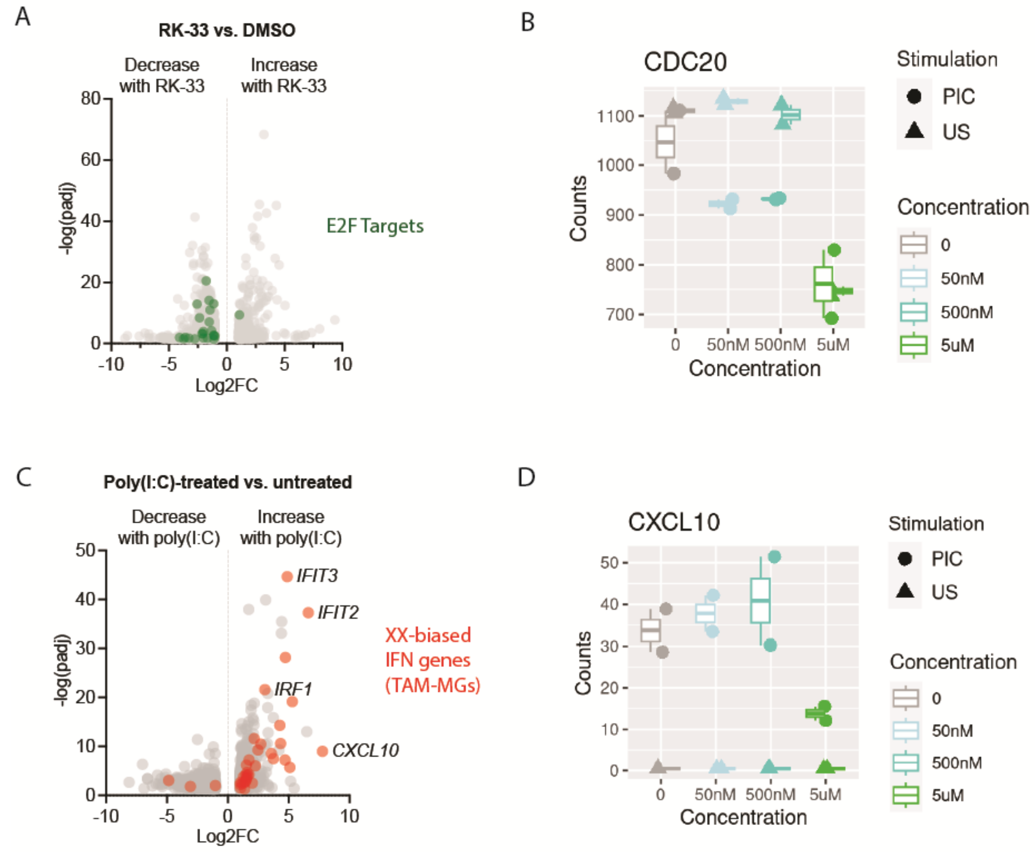


**Supplemental Figure 5.**

A. Genes significantly upregulated and downregulated with 5uM RK-33 treatment in HMC3 microglia. Genes belonging to the E2F targets Hallmark gene set are highlighted in green.

B. Expression of *CDC20* in HMC3 microglia treated with 5uM, 500nM, 50nM RK-33, and DMSO, followed by 10uM poly(I:C) stimulation or unstimulated.

C. Genes significantly upregulated and downregulated with 10uM poly(I:C) treatment in HMC3 microglia. XX-biased IFN alpha genes are highlighted in red.

D. *CXCL10* expression in HMC3 microglia treated with 5uM RK-33 and 10uM poly(I:C).

**Supplemental Tables**

Table S1. TAM-MG and control microglia sample information.

Table S2. TAM-MG and control microglia expressed genes (TPM).

Table S3. TAM-MGs vs control microglia DEGs.

Table S4. Sex-biased genes in low-grade and GBM TAM-MGs.
